# Host cytoskeletal vimentin serves as a structural organizer and an RNA-binding protein regulator to facilitate Zika viral replication

**DOI:** 10.1101/2021.04.25.441301

**Authors:** Yue Zhang, Shuangshuang Zhao, Min Li, Yian Li, Fengping Feng, Jie Cui, Yanhong Xue, Xia Jin, Yaming Jiu

## Abstract

Emerging microbe infections such as Zika virus (ZIKV) pose an increasing threat to human health. Current investigations on ZIKV replication have revealed the construction of replication compartments (RCs) and the utilization of host cellular endomembranes, without careful examination of the cytoskeletal network. Here, we investigated the function of vimentin, one of the intermediate filaments (IFs) that play a central role in basic cellular functions and diseases, in the life cycle of ZIKV infection. Using advanced imaging techniques, we uncovered that vimentin filaments have drastic reorganization upon viral protein synthesis, to form a perinuclear cage-like structure that embraces and concentrates RCs. Genetically removal of vimentin markedly reduced viral genome replication, viral protein production and infectious virions release, without interrupting viral binding and entry. Furthermore, proteomics and transcriptome screens by mass spectrometry and RNA sequencing identified intense interaction and regulation between vimentin and hundreds of endoplasmic reticulum (ER)-resident RNA-binding proteins. Among them, the cytoplasmic-region of ribosome receptor binding protein 1 (RRBP1), an ER transmembrane protein directly binds viral RNA, can interact with vimentin, resulting in modulation of ZIKV replication. Together, our work discovered a dual role for vimentin as being not only a structural element for RCs but also an RNA-binding-regulating hub in the ZIKV infection model, unveiling another layer of the complexity between host and virus interaction.

## INTRODUCTION

Zika virus (ZIKV), a mosquito-borne enveloped RNA virus that belongs to the Flaviviridae family, has gained notoriety recently, due to its explosive outbreaks and association with serious clinical diseases such as Guillain-Barré syndrome in adults and microcephaly in newborns (Cao-Lormeau et al. 2016, Pierson and Graham 2016, Rasmussen et al. 2016, Pierson and Diamond 2018). Currently, no ZIKV-specific therapies or prophylactic vaccines are available (Poland et al. 2019). ZIKV genome is a positive-sense, single-stranded RNA (ssRNA(+)) (Musso and Gubler 2016). The viral replication occurs on the surface of the endoplasmic reticulum (ER), where the double strand RNA (dsRNA) is synthesized from viral genomic ssRNA(+) and transcribed into new proteins (Munjal et al. 2017, Lee et al. 2018). The viral genome is translated into a polyprotein which is proteolytically processed into 3 structural proteins (capsid (C), precursor membrane (prM) and envelop (Env)) and 7 nonstructural proteins (NS1, NS2A, NS2B, NS3, NS4A, NS4B, and NS5), by both host and viral proteases (Shi and Gao 2017, Sirohi and Kuhn 2017).

Vimentin is the most abundant intermediate filaments (IFs) which generally surrounds the nucleus and extends throughout the cytoplasm, providing help to important biological processes such as organelle positioning, cell migration and cell signaling (Lowery et al. 2015). As a highly dynamic filaments that rapidly respond to physiological stimuli through self-assembly and disassembly (Danielsson et al. 2018), vimentin’ role in virus infections has gained increasing attention. For instance, it either acts as a co-receptor to help virus invading target cells, or guides transportation of virus to the replication site, or reorganizes into aggregated structures surrounding replication compartments (RCs), or recruits viral elements to the location of assembly and egress (Dohner and Sodeik 2005, Foo and Chee 2015, Denes et al. 2018, Zhang et al. 2019, Ramos et al. 2020, Zhang et al. 2020). Vimentin network rearrangement has been previously observed in the infection of adenovirus type 2 and type 5 (Defer et al. 1990), African swine fever virus (ASFV) (Stefanovic et al. 2005, Netherton and Wileman 2013), coxsackievirus B3 (Matilainen 2016, Turkki et al. 2020), dengue virus type 2 (DENV-2) (Lei et al. 2013, Teo and Chu 2014), foot-and-mouth disease virus (FMDV) (Gladue et al. 2013, Ma et al. 2020), frog virus 3 (Murti et al. 1988), transmissible gastroenteritis virus (Zhang et al. 2015), vaccinia virus (Risco et al. 2002), and SARS-CoV-2 (Cortese et al. 2020).

A common feature for *Flaviviruses* infection is the induction of cellular endomembrane rearrangements to establish viral RCs, which increase the local concentration of viral and cellular factors for efficient viral replication (Neufeldt et al. 2018, Cortese et al. 2017, Mohd Ropidi et al. 2020). In the case of ZIKV, infection could induce the rearrangement of F-actin in the cell periphery (Nie et al. 2021), and enwrapping of dsRNA-positive structures by bundled microtubules (MTs) and cytokeratin 8 and nestin IFs (Cortese et al. 2017). Perturbation of F-actin by cytochalasin D or Jasplakinolide enhanced ZIKV infection (Nie et al. 2021), while pharmacological treatment with MTs-stabilizing drug paclitaxel strongly reduced ZIKV titer (Cortese et al. 2017). Despite rearrangement of vimentin has been observed in ZIKV infection (Pagani et al. 2017), the dynamic changes of vimentin IFs during ZIKV lifecycle and its contribution to RCs construction and integrity remain understudied.

Besides exploiting cytoskeletal networks, ZIKV can hijack ER-resident proteins, such as ER-localized RNA-binding proteins vigilin and ribosome-binding protein 1 (RRBP1), to facilitate viral genome replication and viral protein translation (Ooi et al. 2019). In addition to provide structural scaffold, there are evidence indicating that cytoskeletal proteins may regulate translational apparatus (Kim and Coulombe 2010). For instance, ribosomes can physically associate with MTs and F-actin in different cells (Hamill et al. 1994, Medalia et al. 2002). Disorganization of F-actin by cytochalasin D impairs local protein synthesis in isolated axoplasmic nerve fibers (Sotelo-Silveira et al. 2008). The interaction between keratin IFs and □ subunit of eukaryotic elongation factor-1 (eEF1B□) plays an essential role in protein synthesis (Kim et al. 2007). MTs can bind to cytoplasmic tail of RRBP1 and take part in ER organization and neuronal polarity (Ogawa-Goto et al. 2007, Farias et al. 2019). However, information on the spatial and functional relationship between vimentin IFs and the translational machinery, especially in the context of virus infection, remain incomplete.

In this study, we investigated the function of vimentin IFs in ZIKV infection. By monitoring spatial-temporal responses of cellular vimentin network throughout various steps of ZIKV life cycle, we demonstrated that ZIKV infection induces massive rearrangements of cytoplasmic vimentin. When vimentin protein was genetically depleted from cells, distribution of viral proteins is scattered within infected cells, and viral RNA replication, protein synthesis, and virion release are subsequently reduced. Using mass spectrometry and RNA sequencing analysis, we discovered interactions between vimentin and RNA-binding proteins, and vimentin binding to RRBP1 facilitates ZIKV RNA replication. Thus, our work establishes important connections among vimentin filaments dynamics, ZIKV RCs, cellular RNA-binding proteins in highly effective infection.

## RESULTS

### ZIKV infection induces drastic cellular vimentin rearrangement and formation of cage-like structures

To study whether host cytoskeletal proteins respond to ZIKV infection, we examined vimentin network using an established model with human osteosarcoma cells (U2OS) that express abundant cytoskeletal filaments and are highly susceptible to viruses (Jiu et al. 2015, Rausch et al. 2017, Hackett et al. 2019) (Fig 1A,E). The cells were infected with an ZIKV Asian lineage strain, SZ01, and fixed at different time points post infection (hpi). Viral structural protein envelop (Env) and endogenous vimentin were visualized by immunofluorescence. In mock infected cells, vimentin filaments showed perinuclear localization and radiated towards cell periphery with apparent filamentous structure (Fig 1A). However, the localization of vimentin rearranged markedly and formed compact aggregation together with viral protein near nucleus, without any notable changes on cell size and overall morphology, within 48 hpi (Fig 1A,B).

**Figure 1.**
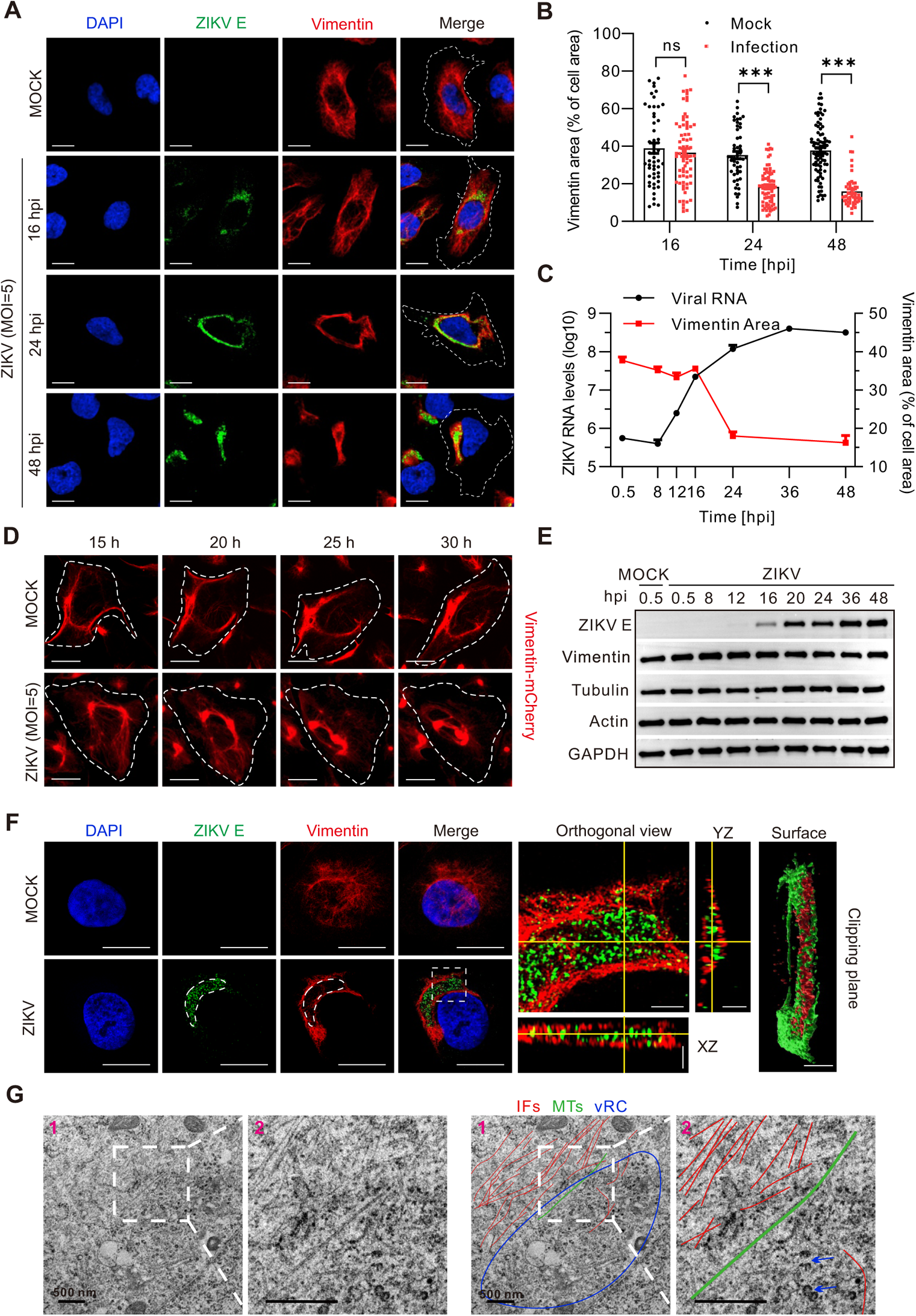
Rearrangement of vimentin filaments in ZIKV-infected cells. **(A)** Human U2OS cells were infected with ZIKV (MOI=5) for 16, 24, and 48 h. Viral envelop (Env) protein and vimentin were stained with respective antibodies, and nuclear was stained with DAPI. The white dotted line indicates the outline of the cell. Scale bar 15 μm. **(B)** Quantification of the proportion of vimentin area versus overall cell area shown in (A). Each point represents a single cell. \*\*\**P*<0.001 (unpaired *t*-test). **(C)** Time course of the accumulated intracellular ZIKV RNA (MOI=0.1) levels measured by qRT-PCR (corresponding to the left axis) and relative vimentin area changes (MOI=5) (corresponding to the right axis), upon ZIKV infection. Error bars indicated means ± SEM from three independent experiments. **(D)** The dynamic rearrangement of vimentin in vimentin-mCherry-expressing cells infected with ZIKV (MOI=5). Scale bar 30 μm. **(E)** Time course of accumulated intracellular ZIKV Env protein level measured by western blot in infected U2OS cells (MOI=0.1). The expression levels of vimentin, tubulin and actin were not affected by ZIKV infection with GAPDH as control. **(F)** Cells were immunostained for vimentin (red) and viral Env (green) at 48 hpi, and visualized by structured illumination microscopy (3D-SIM). White square in the left panel indicates the magnified area shown in the corresponding color image on the right. Orthogonal sections and 3D reconstruction shows that vimentin encapsulates viral Env to form a cage-like structure. Scale bar 15 μm in the left panel, 2 μm in the middle orthogonal view panel, 5 μm in the XZ panel, 5 μm in the YZ panel and 5 μm in the right clipping plane panel. **(G)** Transmission electronic microscopy images of 70 nm thin sections of resin-embedded cells infected with ZIKV (MOI=5). White square indicates the magnified area shown in the corresponding color image on the right. IFs, intermediate filaments indicated with red lines; MTs, microtubules indicated with green lines; vRCs, viral replication compartments indicated with blue circle. Scale bar 500 nm in the cell images.

A time course study showed that viral RNA replication initiated from 8 hpi and reached to a plateau at 36 hpi and onwards, concurrently, viral proteins start to synthetize from 12 hpi and become more pronounced afterwards (Fig 1C,E). It was evident from the quantification that both viral RNA appearance and protein synthesis started before the virus-induced vimentin rearrangement, which was not yet initiated at 16 hpi. The vimentin compartments only appeared to shrink at around 24 hpi, progressively reached to a plateau at 36 hpi (Fig 1B-E). This suggests that vimentin concentration is not required to initiate virus replication. Instead, the emergence of viral proteins may act as a trigger for the vimentin rearrangements to occur.

At the protein levels, however, cytoskeletal components including actin, tubulin and vimentin were similar between ZIKV infected and mock-infected cells throughout the infection lifecycle (Fig 1E). Different from vimentin, neither actin nor tubulin network show significant redistribution during ZIKV infection (Fig S1C), indicating a vimentin-specific response to infection.

Considering vimentin reorganization usually accompanies the assembly and disassembly of the filaments regulated by phosphorylation (Inagaki et al. 1987), which modulates vimentin solubility (Snider and Omary 2014), we performed cellular fractionation experiment and western blot analysis. Immunoblotting showed negligible changes of vimentin content in both the cellular (soluble) and cytoskeletal (insoluble) fraction, as well as the phosphorylation levels from 16 hpi onwards (Fig S1A,B), suggesting that the drastic rearrangement of vimentin was not associated with its assembly turnover or posttranslational modification. To further explore whether microtubules (MTs) are the machinery responsible for vimentin transportation towards nucleus, we applied two drugs inhibiting assembly of MTs and the activity of its retro-wards motor dynein, respectively, and found that these drugs did not prevent virus induced vimentin retraction, ruling out the possible involvement of MT-dominated aggresomal transportation system (Fig S1D).

To describe the detailed dynamics of vimentin filaments rearrangement, we generated a vimentin-mCherry stable expression cell line and performed live-cell imaging analysis. A high MOI of input virus was used to achieve infection of every cell recorded. In mock-infected cells, no apparent changes of vimentin were observed within a 30 h monitoring period (Fig 1D). In contrast, vimentin filaments gradually gathered around nucleus from ~20 hpi and progressively intensified in ZIKV-infected cells (Fig 1D; Video S1,S2).

ZIKV multiplies in perinuclear replication compartments (RCs) (Welsch et al. 2009; Caldas et al. 2020). In order to gain insight into the spatial relationship between vimentin and viral protein, we took advantage of the three-dimensional-structured illumination microscopy (3D-SIM) for super-resolution visualization. Side view of 3D images discovered that vimentin filaments form a hollow cage-like structure that wrap structural protein Env as well as nonstructural proteins NS1 and NS4B (Fig 1F; Fig S1E). Furthermore, electronic microscopy elaborated events of enrichment of IFs-like filaments next to the concentrated area of viral particles in the perinuclear region (Fig 1G; Fig S1G). Together, these imaging observations indicated that ZIKV infection induced vimentin-cage formation around RCs.

### Exogenous expression of Zika viral protein leads to vimentin enrichment in the perinuclear area

To further dissect the role of viral protein in vimentin rearrangement, we transfected vimentin-mCherry-expressing cells with moderate concentrations of EGFP tagged Env protein and monitored vimentin dynamics. The results showed that during 1.5 h real-time monitoring after overnight transfection, vimentin filaments and viral Env protein accumulated next to nucleus synchronously (Fig 2A; Video S3). By tracing the fluorescent intensity of perinuclear region, vimentin was found to be gathered significantly faster (Fig 2B).

**Figure 2.**
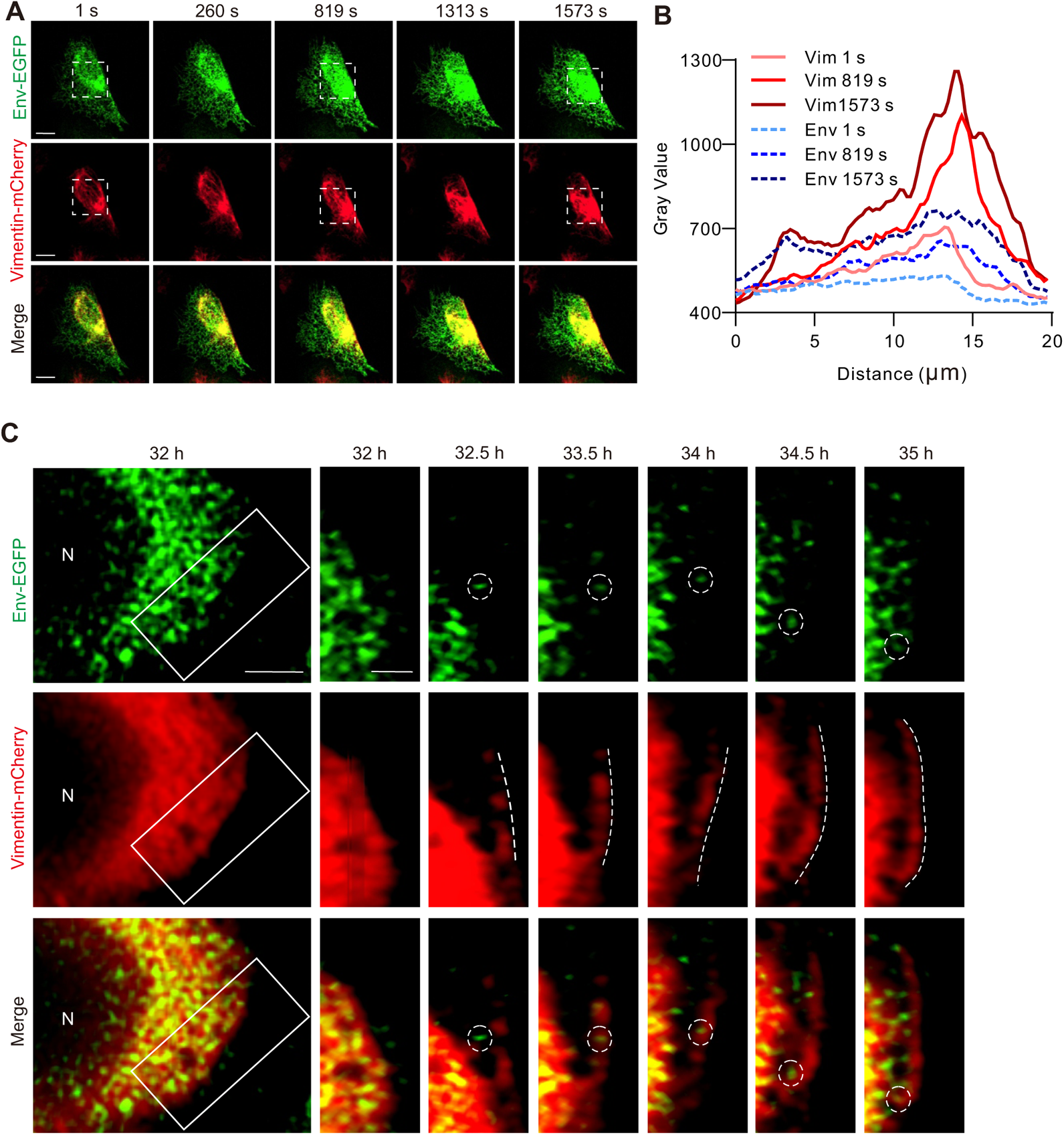
Expression of ZIKV envelop protein induces vimentin rearrangement. **(A)** The dynamic rearrangement of vimentin in vimentin-mCherry-expressing cells transfected with ZIKV Env-EGFP plasmid. Scale bar 10 μm. **(B)** Plot profile analysis of Env-EGFP and vimentin-mCherry intensities in the white doted box in (A) at corresponding time point. Distance in Axis X represents horizontal distance through the selection. **(C)** Vimentin filaments gather scattered viral Env protein towards peri-nucleus region. N and white box in the left panels indicate cell nucleus and the magnified area shown in the corresponding right panel. The dotted circles indicated the positions of one ZIKV-Env foci, and the dotted white lines indicated the shape of one vimentin filament. Scale bar 2 μm in the cell images and 1 μm in the magnified images.

Moreover, the retrograde movement of vimentin filaments gather the scattered viral protein, as visualized by individual fluorescent foci, to the perinucleus region (Fig 2C; Video S4), indicating a potential for vimentin being a pro-viral factor. Together, these results suggest that the cellular machinery recognizing foreign viral protein triggers events leading to varied vimentin arrangement.

### Host vimentin is required for the integrity of viral replication compartments and efficient infection

Vimentin has gained more attention on its roles in various viral infections (Dohner and Sodeik 2005, Foo and Chee 2015, Denes et al. 2018, Zhang et al. 2019, Ramos et al. 2020, Wen et al. 2020, Zhang et al. 2020). To determine whether there is a causal relationship between vimentin rearrangement and ZIKV infection, we used CRISPR/Cas9 method to establish vimentin-knockout (KO) in U2OS cells and Huh7 (Fig S2A,D). Results showed that reducing vimentin levels neither affected the cell viability nor cell growth (Fig S2B,C,E). We next used lentiviral system to establish vimentin KO-full length (FL) rescue cells, and verified the levels of vimentin protein express in these cells by western blot (Fig S2A).

In these cells, we observed not only similar vimentin network shrinkage and formation of a concrete compartment for viral RCs, but also viral protein localization at a subcellular level (Fig 3A). Intriguingly, viral protein staining for Env, NS1 and NS4B displayed a piecemeal distribution in vimentin KO cells (Fig 3A, Fig S2I,J), but rarely seen in wild-type cells. Moreover, depletion of vimentin dramatically increases the proportion of partitioned cellular viral protein compartments up to 85% and reduces the total viral protein area, both of which can be fully rescued when full-length vimentin was reintroduced (Fig 3A-C). Nevertheless, vimentin which forms unit length filaments (ULF) or squiggles was not able to fully rescue the viral dispersion phenotype observed in infected cells (Fig 3A,B).

**Figure 3.**
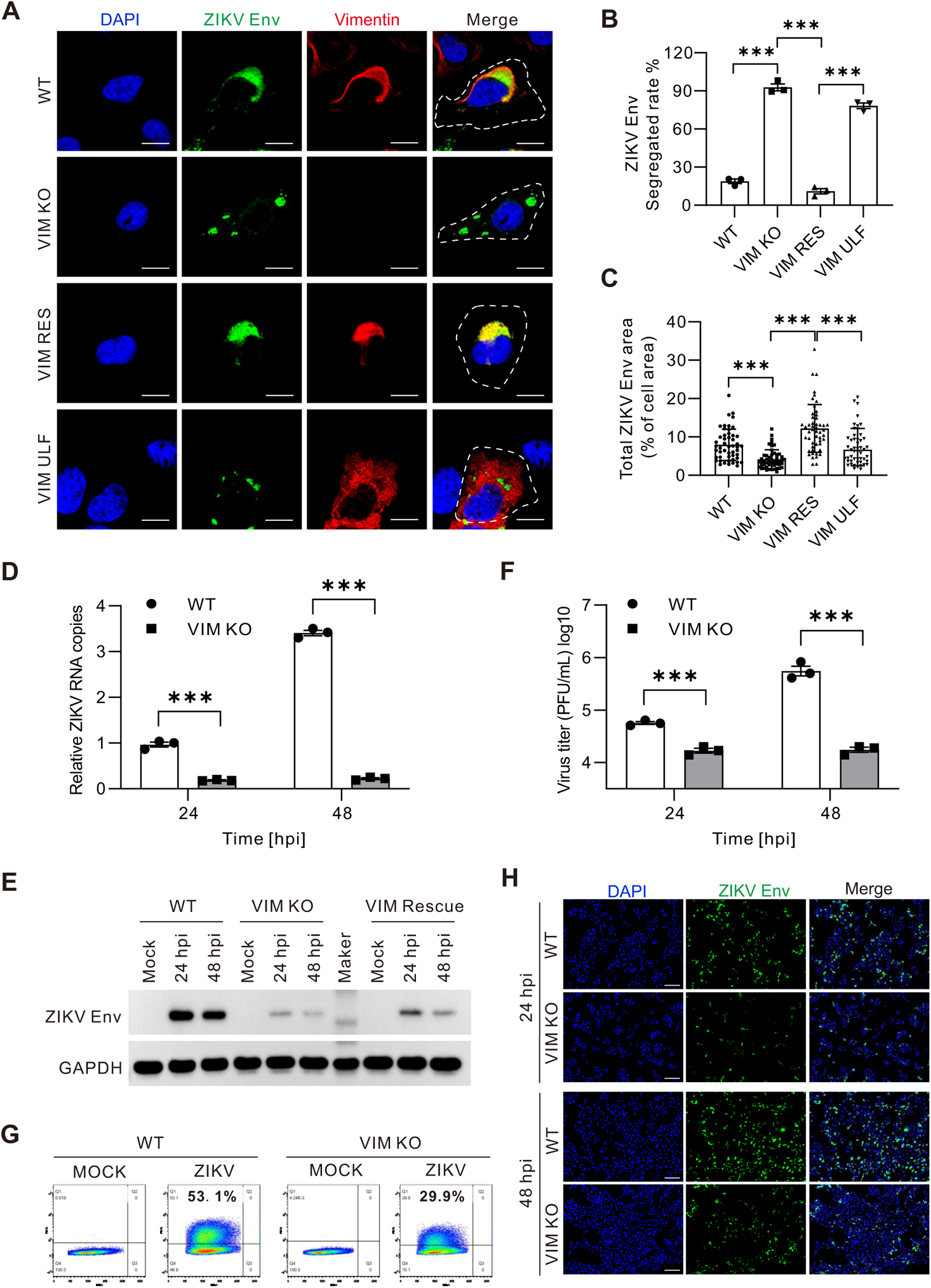
Depletion of vimentin results in disruptive replication compartments and subsequent reduced ZIKV infection. **(A)** WT, vimentin KO (VIM KO), vimentin-full-length re-introducing (VIM RES) and vimentin-unit-length-filament (VIM ULF) re-introducing cells were infected with ZIKV (MOI=5) for 24 h. Cell were fixed and immunostained to visualize ZIKV Env, vimentin and nucleus. Scale bar 15 μm. **(B)** Quantification of the percentages of cells with segregated ZIKV Env in WT, VIM KO, VIM RES and VIM ULF conditions. Each point represents an independent experiment. **(C)** Quantification of the percentage of total ZIKV Env area to overall cell area in WT, VIM KO, VIM RES and VIM ULF conditions. Each point represents an infected cell. **(D-F)** Intracellular ZIKV RNA (D), ZIKV Env protein level (E) and titers of ZIKV particles (F) in infected WT and vimentin KO cells (MOI=0.1) were measured by two-step qRT-PCR, western blotting, and plaque assay, respectively. Results from three independent experiments are shown. **(G)** Percentage of infected WT and VIM KO cells (MOI=1, 48 hpi) measured by flow cytometry. **(H)** Percentage of infected WT and VIM KO cells (MOI=5) measured by immunofluorescence. Scale bar 100 μm. Error bars indicated means ± SEM.

How this scattered subcellular distribution of viral protein influences the infectivity? To address this, we subsequently measured viral genome replication by qRT-PCR and viral protein expression by western blot in vimentin KO cells. It is apparent that the absence of vimentin caused significant reduction of viral RNA copies and structural protein Env levels at both 24 and 48 hpi (Fig 3D,E), and reintroducing vimentin partially rescued the viral protein expression (Fig 3E). Virus titer was further measured and as expected, there was at least 20-fold less of viruses released from vimentin depleted cells at 48 hpi (Fig 3F). Similarly, in Huh7 cells, vimentin depletion showed defective infection as that in U2OS cells (Fig S2F-H).

By immunostaining and flow cytometry, fewer infected cells were detected in vimentin depletion background at 24 and 48 hpi, the fast replication period during viral life cycle (Fig 3G,H), suggesting that vimentin is critical for efficient ZIKV infection. Taken together, depletion of host vimentin leads to compromised viral infection as demonstrated by less genome replication, reduced protein expression, and fewer particle production, most probably as a result of the disruption of the integrity of concentrated perinucleus RCs.

### Vimentin depletion compromises viral genome and protein synthesis without affecting viral binding and entry to the host cell

To further investigate the function of vimentin, a time course of infection experiment was performed. Changes of viral RNA copies in wild-type and vimentin KO cells were confirmed by qRT-PCR. Although the viral RNA numbers were the same at the beginning of infection, their copies in vimentin KO cells were significantly lower than that in wild-type cells during the later period of infection (Fig 4A). Apart from the replication dynamics of viral genome, viral proteins in both backgrounds accumulated gradually, but vimentin depletion led to a slower accumulation and thus much less viral Env protein expression at each time point from 16 hpi onwards (eg. 20, 24, 36, 48 hpi), compared to wild-type cells (Fig 4B).

**Figure 4.**
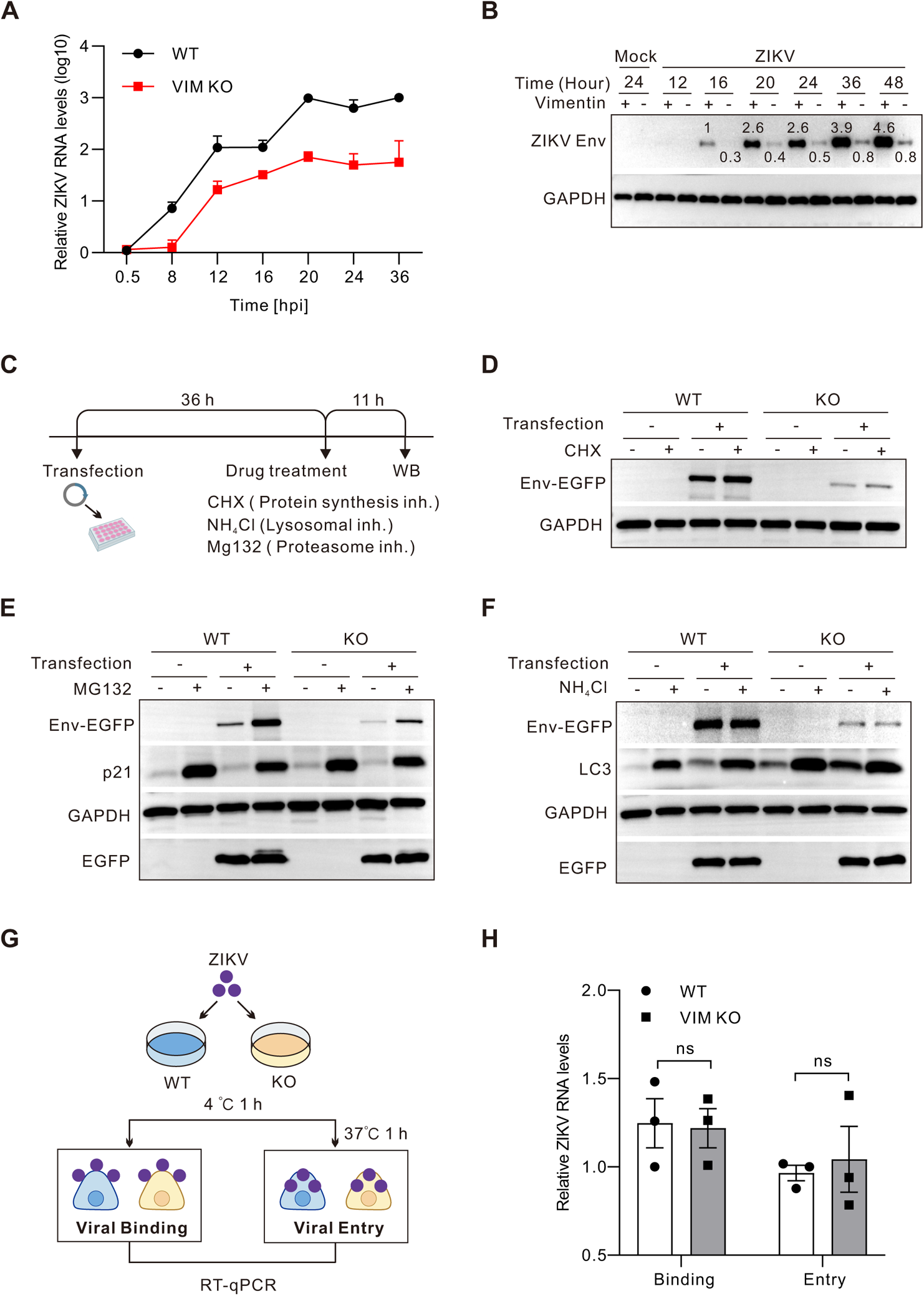
Depletion of vimentin influences the production and stability of viral components after entering cells. **(A-B)** Time course of accumulated intracellular ZIKV RNA in infected WT and vimentin KO (VIM KO) cells (MOI=0.1) measured by qRT-PCR. **(B)** Time course of accumulated intracellular ZIKV Env protein in infected WT and VIM KO cells (MOI=0.1) measured by western blots. Numbers in the blots indicated the levels of ZIKV Env normalized to GAPDH. **(C)** The schematic diagram of drug treatment experiment. Cells were transiently transfected with E-EGFP plasmid for 36h, and then treated with MG132 (10 μM), NH_4_Cl (25 mM) and CHX (20 μg/mL) for 11 h. **(D)** Western blotting analysis of viral protein upon CHX treatment. **(E)** Western blotting analysis of viral protein upon MG132 treatment, where detection of P21 accumulation serves as positive control. **(F)** Western blotting analysis of viral protein upon NH_4_Cl treatment, where detection of LC3 accumulation serves as positive control. **(G)** The schematic diagram of ZIKV binding and entry assay. **(H)** ZIKV RNA levels from bound and internalized ZIKV were measured by qRT-PCR. The data are from three independent experiments. ***P<0.001 (2way ANOVA). Error bars indicated means ± SEM.

The observed low levels of viral Env protein expression in vimentin KO cells could be due to a decrease in protein biogenesis and/or an increase in protein degradation. We therefore tested whether vimentin could influence the stability of ZIKV proteins. Plasmid expressing Env-EGFP fusion protein was transiently transfected into cells and analyzed by western blot 48 h later. The results showed that vimentin depletion dramatically reduced the exogenous expression of Env-EGFP (Fig 4D-F). We next treated cells with cycloheximide (CHX, 20 μg/mL for 11 h) to inhibit protein translation, and found that viral Env was relatively stable in both wild-type and vimentin KO cells (Fig 4D). These suggest that vimentin acts on protein biosynthesis but not degradation. The following experiments further confirmed this assertion.

There are two classic destinations for cellular protein degradation regulated by proteasomal and lysosomal pathways. To further analyze whether depletion of vimentin promotes the degradation of viral protein and via which way, cells transfected with Env-EGFP were treated with proteasomal specific inhibitor MG132 (10 µM for 11 h), or lysosomal specific inhibitor ammonium chloride (NH4Cl) (25 mM for 11 h) (Fig 4C). Immunoblot analysis showed that the Env-EGFP protein level apparently increased in both wild-type and vimentin depletion cells after MG132 treatment (Fig 4E), but not influenced by NH4Cl application (Fig 4F). Concurrently, p21 and LC3, known proteins for proteasome and lysosomal degradation, respectively, were chosen as positive controls for these experiments. Combing results of the increased Env upon MG132 treatment and unaltered Env upon NH4Cl treatment, it is reasonable to conclude that Env protein was degraded mainly via proteasomal pathway and vimentin depletion does not interrupt the degradation of Env protein.

Given that vimentin shrinking was visualized from 24 hpi but not before, we speculated that the early steps of viral replication were not affected in vimentin depletion cells. To confirm this, we used two assays to examine binding and entry steps (Fig 4H), respectively. Cells were infected by ZIKV for 1 h at 4□, directly or incubated for 1 more hour at 37 □, washed and then cell lysates collected for measurement of RNA copies by RT-qPCR (Fig 4G). The results confirmed that both the binding and entry of ZIKV are similarly efficient in wild-type and vimentin KO cells (Fig 4H). Collectively, these results suggest that vimentin acts on the production and accumulation of ZIKV proteins as well as the efficient replication of ZIKV RNA, without influencing viral internalization.

### A large number of host RNA-binding proteins interact with and being regulated by vimentin during ZIKV infection

Since the perinuclear expression of vimentin is highly associated with ER where viral RNA replication and protein translation take place, we speculate that aside from being a structural scaffold, vimentin may interact with host factors that regulate viral RNA transcription and subsequent protein synthesis. To test this, we implemented mass spectrometry (MS) analysis to identify proteins interacting with vimentin, and subjected 1050 candidates to gene ontology (GO) analyses by DAVID (The Database for annotation, visualization and integrated discovery). The recognized vimentin interactors were classified based on three taxonomic features including molecular function, cellular component and biological process. From the classification analysis, we found that a large proportion of candidates interacted with vimentin are RNA binding proteins and ribosome components (Fig 5A), and they are intimately related to RNA processing, translation and viral transcription (Fig 5B). Importantly, GO annotation revealed a significant enrichment of ER components with vimentin (Fig 5C), indicating that vimentin is the principal factor that interacts with ER-associated RNA-binding proteins (RBPs) in host cells.

**Figure 5.**
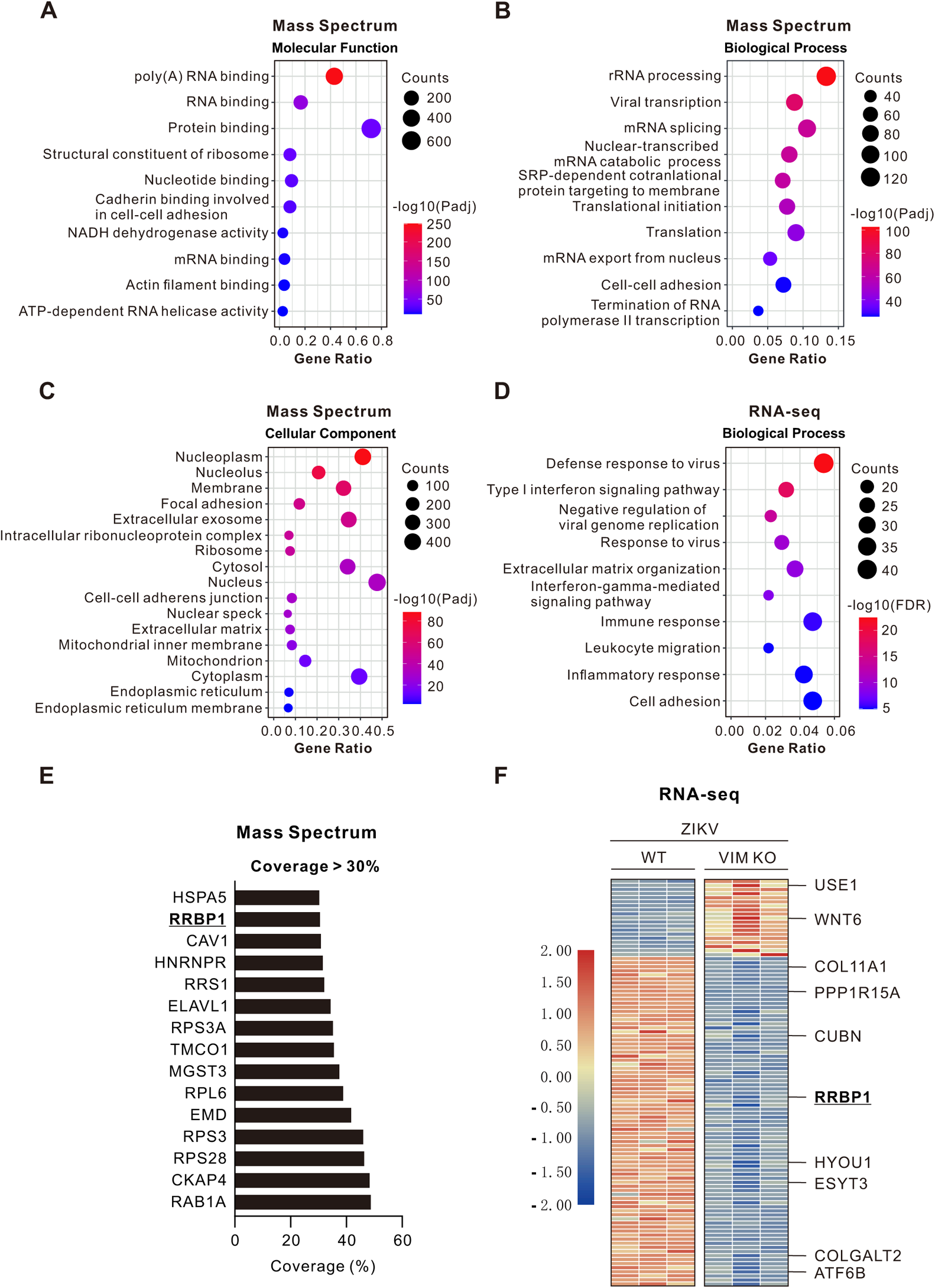
Interaction between vimentin and ER-localized RNA-binding components affects ZIKV infection. **(A)** The top 10 significant GO terms in molecular function are shown in the bubble chart. **(B)** The top 10 significant GO terms in biological process are shown in the bubble chart. **(C)** All 17 significant cellular components with count number greater than 50 genes are shown in the bubble chart. **(D)** Biological process enrichment analysis of significantly down-regulated genes in ZIKV infected VIM KO cells compared with infected WT cells (MOI=1, 24 hpi) by RNA-Seq. The top 10 enriched terms are shown in the bubble chart. In (A-D), the color of the bubbles displayed from red to blue indicated the descending order of −log10(Padj). The sizes of the bubbles are displayed from small to large in ascending order of gene counts. The x and y axis represent the gene ratio and the GO terms, respectively. **(E)** List of ER-located proteins interacting with vimentin by setting the coverage peptides threshold over 30% from mass spectrometry assay. **(F)** Heatmap of significantly differentially expressed genes (ER-annotated) between ZIKV infected WT and VIM KO cells (MOI=1, 24 hpi) identified by RNA-Seq (n=3 independent experiments per condition).

To complement with the interactome assay, RNA sequencing (RNA-Seq) was subsequently applied to analyze the variations of global gene transcription in both wild-type and vimentin KO cells infected (or not) with ZIKV. A total of 1408 genes were significantly affected (*P*<0.05), including 518 up- and 890 down-regulated genes in vimentin depletion background upon ZIKV infection (Fig S3C). Among the downregulated hits, there are many ER-localized genes related to double-strand RNA binding and transcription regulation (Fig S3A,B), as well as a large proportion of genes involved in antiviral immune and inflammatory response (Fig 5D). Together with the MS result, we conclude that aside from contributing to RCs scaffolding which provides a structural support, vimentin is also involved in processing viral RNA replication and serving a functional role.

Given the critical role of ER in ZIKV infection, we more carefully examined candidates with ER localization in both MS and RNA-Seq results (Fig 5E,F). By investigating ER annotated candidates, we obtained top 15 vimentin-interacting proteins by setting the threshold of interacting peptides coverage over 30% from MS results (Fig 5E; Table S1). Concurrently, RNA-Seq data were analyzed and the results demonstrated dramatical changes of the mRNA levels of many ER-associated genes, among them, 80% of significantly differentially expressed genes were downregulated (Fig 5F). By crosschecking MS and RNA-Seq candidates, ER-resident protein ribosome-binding protein 1 (RRBP1, also known as p180) was recognized as the only common hit, which shows high endogenous expression in wild-type and substantially down-regulated expression in vimentin KO cells during ZIKV infection (Fig 5F). Of note, it has been recently reported that RRBP1 play a pronounced role in flaviviruses (eg. DENV, ZIKV) infection by directly binding to viral RNA (Ooi et al. 2019). We thus focused on RRBP1, and asked how the interaction between vimentin and RRBP1 influence ZIKV infection.

### ER-resident RRBP1 interacts with and regulated by vimentin to catalyze ZIKV infection

RRBP1 contains a short ER luminal domain, a transmembrane domain, and a large cytoplasmic domain containing tandem repeats motif (Reid and Nicchitta 2015) (Fig 6A). RRBP1 peptides interacting with vimentin identified from MS were located mostly in the cytoplasmic coiled-coil region (25 out of 28 recognized peptides) (Fig 6A, Table S2). We subsequently performed the pull-down assay by using purified His-tagged vimentin protein with an RRBP1 antibody as probe. The results confirmed that RRBP1 indeed interacts with vimentin both *in vivo* and *in vitro* (Fig 6B).

**Figure 6.**
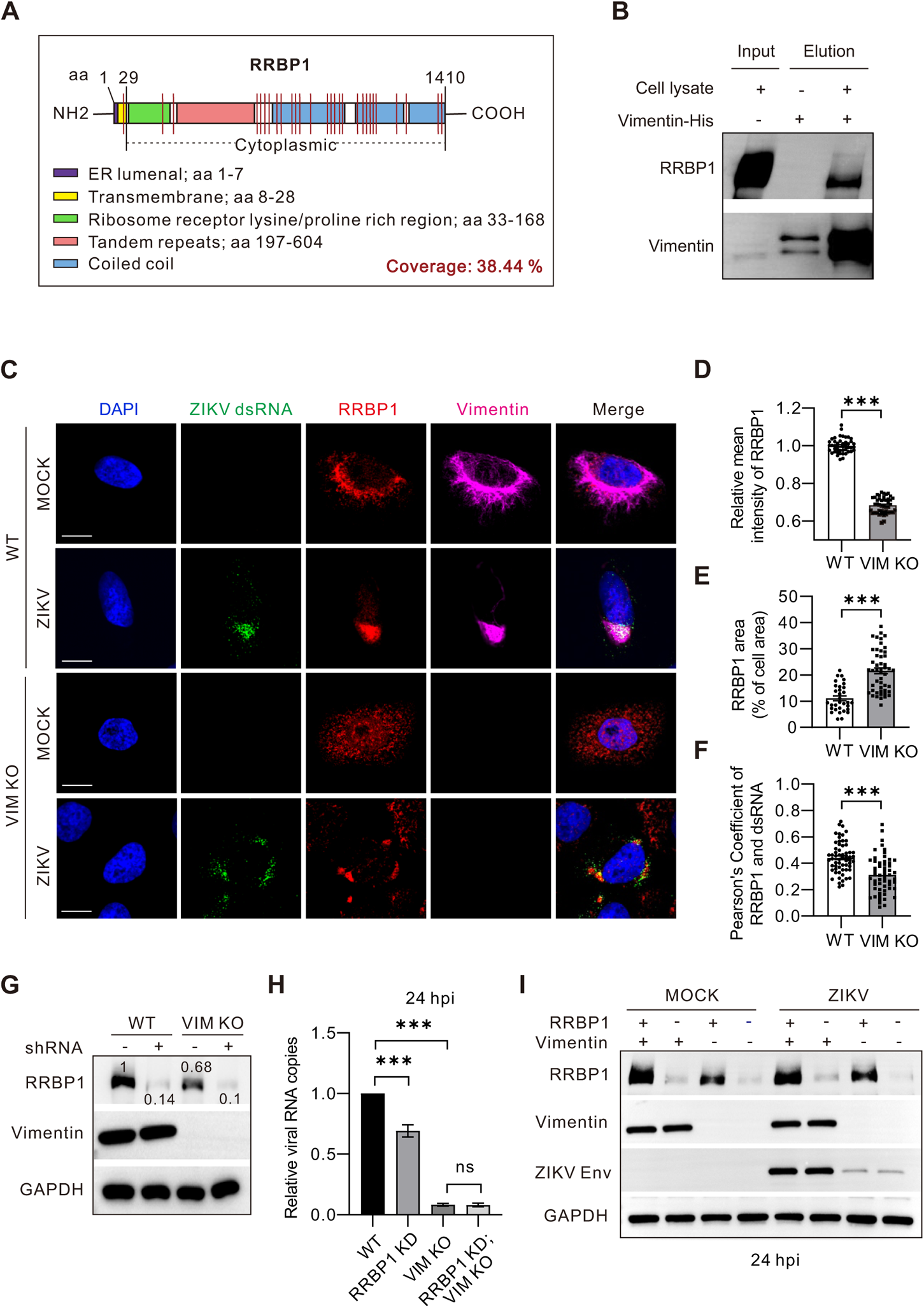
Vimentin interacts with ER-located RRBP1 to regulate ZIKV infection. **(A)** The domain structure of RRBP1 protein. Red lines represent the peptides interacting with vimentin identified from mass spectrometry. **(B)** *in vitro* binding assay of vimentin with RRBP1. Cell lysates were incubated with purified recombinant his-tagged vimentin protein, and analyzed by anti-RRBP1 antibody in western blotting. **(C)** Immunofluorescence images of vimentin, RRBP1 and ZIKV RNA in WT and vimentin KO (VIM KO) cells infected with ZIKV (MOI=2) for 24 h, and stained with anti-dsRNA, anti-RRBP1 and anti-vimentin antibodies and DAPI for nucleus. Scale bar 15 μm. **(D,E)** Quantifications of RRBP1 intensities (D) and areas (E) in non-infected (MOCK) WT and VIM KO cells. Each point represents a single cell. \*\*\**P*<0.001 (unpaired *t*-test). **(F)** Quantification of the colocalization between RRBP1 and dsRNA in ZIKV infected WT and VIM KO cells by Pearson’s coefficients. Each point represents a single cell. \*\*\**P*<0.001 (unpaired *t*-test). **(G)** Western blots demonstrated that RRBP1 was efficiently knockdown by shRNA in both WT and VIM KO cells. Numbers in the blots indicated the levels of RRBP1 normalized to GAPDH. **(H)** Quantifications of ZIKV RNA copies detected by qRT-PCR, and normalized to GAPDH, in WT, RRBP1 knockdown (RRBP1 KD), VIM KO and RRBP1 knockdown; vimentin knockout (RRBP1 KD;VIM KO) conditions. The cells were infected with ZIKV at MOI=0.1 for 24 h, and the data are from three independent experiments. **(I)** Protein levels of RRBP1, vimentin, ZIKV Env in WT and vimentin KO cells detected by western blots, which were also probed with vimentin antibody to confirm the KO efficiency and GAPDH antibody to verify equal sample loading.

Next, we explored the correlation between the cellular expression of vimentin and RRBP1 upon ZIKV infection. Immunofluorescence results showed that in wild-type cells, both vimentin and RRBP1 are accumulated around nucleus where ER network resides (Fig 6C). Upon ZIKV infection, both vimentin and RRBP1 aggregated near the nucleus and co-localized with dsRNA-staining positive viral RNA (Fig 6C; Fig S4D), indicating both of them have participated in the viral RCs process.

To elucidate the relationship between vimentin and RRBP1, we compared the RNA level, protein level and the localization of RRBP1 in wild-type and vimentin KO cells. Results showed that depletion of vimentin reduced both the mRNA and protein levels of RRBP1 to around 50%, and enlarged the cellular distribution of RRBP1 (Fig 6D,E; Fig S4A-C,E). In contrast, deprivation of RRBP1 neither affect the mRNA nor the protein level of vimentin (Fig 6G; Fig S4F). Imaging data showed that lacking vimentin led to an expansion of the subcellular RRBP1 in uninfected cells (Fig 6C). Consistently, upon ZIKV infection, RRBP1 turned into scattered segregation in vimentin KO cells, similar as segregated viral proteins (Fig 3A; Fig 6C), indicating that disruption of RCs integrity by vimentin influenced not only viral components but also host viral-binding components. It is subsequently noted that the colocalization between RRBP1 and viral dsRNA were significantly reduced (Fig 6C,F), suggesting that vimentin depletion reduced the combined abundance of viral-host constituents in RCs. Therefore, these results suggested that RRBP1 is one of the effectors of vimentin in both normal and infection conditions, through which ZIKV replication was affected.

To verify this, we generated RRBP1 knockdown (KD) cells in both wild-type and vimentin KO background by shRNA (Fig 6G; Fig S4E,F) and then infected them with ZIKV. Consistent with a previous study (Ooi et al. 2019), RRBP1 KD reduced viral RNA copies by about 40% (Fig 6H). In comparison, the effect of vimentin depletion on the reduction of viral genome replication was much more severe than that of RRBP1 KD; however, the double depletion has no additive effects (Fig 6H), suggesting that RRBP1 and vimentin located in the same regulating cascade during ZIKV infection. Next, synthesized viral protein was analyzed. The absence of vimentin resulted in significant reduction of structural protein Env, but the absence of RRBP1 alone showed no obvious effect (Fig 6I), indicating that RRBP1 plays a role in viral RNA replication rather than protein synthesis. Taken together, these results demonstrate a pivotal role of vimentin as an upstream factor regulating one of the known viral-binding host factor RRBP1 to affect viral RNA replication. Combined with omics analysis, these data further suggest that aside from contributing to scaffolding RCs, vimentin also acts as a hub molecule to interact with and regulate various RBPs. Both functions of vimentin are important for facilitating efficient ZIKV replication and promoting infection.

## DISCUSSION

In this study, we report the spatial-temporal rearrangement of vimentin filaments induced by ZIKV infection. Importantly, we reveal two major functions of host cell vimentin during ZIKV infection: (1) ‘organizer’, as a structural support to increase the local concentration of all necessary factors for high efficiency of viral replication; (2) ‘regulator’, by interacting with and regulating RNA-binding proteins, such as RRBP1, to facilitate viral replication. The latter is the first time that vimentin has been demonstrated to play a non-scaffold function in the context of virus infection.

### Vimentin cage formation in viral infections

Viruses replicate more efficiently within the viral replication compartments (RCs), but the structural features of such a compartment remain elusive. Our current study uses ZIKV to examine this issue. ZIKV infection induces nestin, one of the intermediate filaments (IFs) in neural stem and progenitor cells, to restructure in the perinuclear region to wrap around viral dsRNA (Cortese et al. 2017). Whether vimentin IFs that expressed in most cell types with greater abundance have a role in ZIKV infection have not been explored. Taking advantage of imaging techniques, we found that ZIKV infection led to vimentin remodeling and formation of cage-like structures that surround the RCs, long after the initiation of viral RNA replication, but following more closely to viral protein production (Fig 1C,E). This kinetic association between host vimentin and viral products has been demonstrated by several lines of evidence: (1) the appearance of vimentin shrinking was approximately 16 h from the commencement of infection, coincides with the period of exponential increase of viral RNA replication (Fig 1C). (2) vimentin filaments gradually accumulated to the site of viral protein synthesis or RCs, to eventually form a whole cage (Fig 1A). (3) the cages could also be formed when cells started to synthesis new viral proteins (Fig 2A,C). Finally, (4) cage formation was not observed in cases where only EGFP vectors were expressed, or medium without live viruses was included (Fig S1F). Despite unable to prove a cause-and -effect mechanism as yet, the above data strongly suggest a link between ZIKV infection and vimentin function.

In agreement with our data, vimentin rearrangement has been observed in other experimental models of infection. In cells with DENV-2 infection, vimentin has increased the interaction with viral NS4A protein, and vimentin filaments gradually moved towards the nucleus, and finally formed cage structures at around 48 hpi (Teo and Chu 2014). In SARS-CoV-2 infection, the beginning of vimentin retraction falls within the timeframe (around 6.5 hpi) of genome replication (Cortese et al. 2020). Both the time of appearance and the co-localization in the perinuclear area for the replication-assembly organelles, leading us to interpret that vimentin sensed viral replication and formed a scaffold to facilitate viral replication.

### The underlying mechanisms of vimentin cage formation in viral infections

Several mechanisms have been postulated for vimentin rearrangement during viral infection. One suggests that vimentin rearrangement is due to its phosphorylation on specific sites by viral infection-induced kinase activation. For example, DENV-2 infection can induce Rho-associated protein kinase (ROCK) activation and phosphorylation of vimentin at serine 72 (Ser72) (Lei et al. 2013), as well as calmodulin-dependent protein kinase II (CaMKII) activation and phosphorylation of vimentin at Ser39 (Teo and Chu 2014). AFSV replication results in the activation of CaMKII and phosphorylation of vimentin at Ser83 (Stefanovic et al. 2005, Netherton and Wileman 2013). In contrast, we found the phosphorylation levels of vimentin at Ser39, Ser56, Ser83 as well as the soluble/insoluble ratio of vimentin have no significantly fluctuation during ZIKV infection (Fig S1A,B), indicating the phosphorylation mechanism for vimentin reorganization may vary among viruses, and ZIKV induced vimentin remodeling is not through phosphorylation at least on these sites nor the solubility regulation.

Aside from phosphorylation, crosslinking of neighboring vimentin subunits through cysteine residue at cysteine 328 (Cys328) in response to oxidative stress can result in reorganization of vimentin network (Perez-Sala et al. 2015). Whether vimentin Cys328 plays a similar role in respond to massive virus replication remains uncertain (Ledur et al. 2020).

Another mechanism is related to the aggresome processing machinery including dynein, dynactin, and microtubules (MTs). For instance, the capsids of influenza A virus (IAV), with the help of aggresome processing machinery, can mimic misfolded protein aggregates and start uncoating in the cytoplasm (Banerjee et al. 2014). Inhibition of dynein-dependent transport by overexpression of p50 (also known as dynamitin) block the recruitment of vimentin to the microtubule organizing center (MTOC) during ASFV infection (Stefanovic et al. 2005). Nevertheless, we found that neither chemical inhibition of dynein by cilibrevin D nor inhibition of MTs by nocodazole has influenced the perinuclear aggregation of vimentin and Env protein during ZIKV infection (Fig S1D).

Overall, our results exclude the above mechanisms of vimentin reorganization during ZIKV infection, negating the role of phosphorylation modification or aggresomal machinery. Therefore, further work is needed to unravel the molecular mechanisms of vimentin filaments rearrangement in ZIKV infections.

Finally, direct interaction between viral protein and cellular vimentin may also trigger the formation of vimentin cage. For instance, human immunodeficiency virus 1 (HIV-1) protease can cleave vimentin bundle and then induce accumulated perinuclear localization of vimentin filaments (Shoeman et al.1990, Shoeman et al. 2001, Honer et al. 1991). In the case of DENV-2 and FMDV infections, vimentin interacts with nonstructural protein DENV-2 NS4A and FMDV 2C, respectively, concurrent with the formation of vimentin cage (Gladue et al. 2013; Teo and chu 2014). Our results are more consistent with the interpretation that ZIKV proteins may directly or indirectly interact with vimentin, which cooperates with other host factors, causing the formation of vimentin cage.

### The need of vimentin in viral infections

In line with the hypothesis that the cytoskeleton cage observed in ZIKV infection might contribute to spatially concentration of different viral-induced membranous structures (Cortese et al. 2017), we demonstrated that without the cage formation after vimentin depletion, RCs are miss-organized and segregated in the cytoplasm (Fig 3A), leading to less efficient synthesis of viral components, and lower overall infection efficiency (Fig 3A,D-H; Fig 4A,B). In DENV-2 infection, the RCs become diffused throughout the cytoplasm when vimentin was knockdown by siRNA (Teo and Chu 2014). Differently, we witnessed that ZIKV RCs are scattered into lumps rather than evenly diffused as in DENV-2 case, indicating a distinct dispersion feature regulated by vimentin upon ZIKV infection. Moreover, disruption of vimentin filaments by a drug Acrylamide significantly reduced the release of both bluetongue virus (BTV) and DENV-2 (Bhattacharya et al. 2007, Kanlaya et al. 2010). Treatment with Withaferin A, a compound that disrupts vimentin network, resulted in a significant reduction in SARS-CoV-2 replication and virion released (Cortese et al. 2020). Thus, it may be a common function in viral infection that vimentin cage organizes the replication structures and provides an optimal niche for the replication/translation to occur.

The decreased viral protein level within cells may be due to the reduced production or increased degradation. Vimentin has been previously shown to regulate the proteasomal degradation of HCV core protein to affect HCV production (Nitahara-Kasahara et al. 2009). In contrast, by treatment with protein synthesis and degradation inhibitors, we showed that ZIKV Env biogenesis, but not its stability, was reduced in vimentin depleted cells (Fig 4C-F). Thus, why vimentin acts differently in different viral infections remain to be determined.

### The non-structural role of vimentin in viral infections

The ER is an essential cellular compartment for completion of the virus life cycle. During ZIKV replication, there is an accumulation of viral components in the ER (Mohd Ropidi et al. 2020), and viral nonstructural proteins can be incorporated into ER membrane to create invagination or protrusion vesicles for viral RNA replication (Neufeldt et al. 2018).

Since viruses cannot encode all proteins necessary for their life cycle, they usurp cellular protein biogenesis machineries such as ribosomal proteins (Campos et al. 2017), RNA-binding proteins (RBPs) for viral RNA replication/transcription (Garcia-Moreno et al. 2018, Dicker et al. 2020, Diosa-Toro et al. 2020), and formation of ER membrane protein complex (EMC) (Barrows et al. 2019, Lin et al. 2019). Noncoding subgenomic flavivirus RNA (sfRNA) produced by ZIKV can interact with over 20 RBPs to regulate multiple cellular post-transcriptional processes and therefore limit effective response of these cells to viral infection (Michalski et al. 2019, Jansen et al. 2021). 464 RBPs was identified being associated with DENV or ZIKV gRNAs, including previously reported candidates (eg. heterogeneous nuclear ribonucleoproteins (hnRNPs), polyadenylate-binding protein (PABP)) that specifically associate with DENV RNA (Phillips et al. 2016, Viktorovskaya et al. 2016, Ooi et al. 2019, Scaturro et al. 2019), and recently known RBPs vigilin and RRBP1 which were reported to directly bind to DENV and ZIKV RNA (Ooi et al. 2019).

In line with these discoveries, using mass spectrometry and RNA sequencing analysis, we revealed that vimentin interacts with a large number of RBPs and ribosomal proteins to regulate cellular transcription and translation, enabling efficient ZIKV replication (Fig 5A-F). Considering the interplay between cytoskeleton and ER membrane (Terasaki et al. 1986, Risco et al. 2002, Gurel et al. 2014, Zhang 2020), it is tempting to speculate that virus-induced vimentin cage not only provides physical space for viral RCs accumulation, but equally important, interacts with molecules involved in cellular transcription and translation process and thus promotes the efficiency of virus replication from perspectives of both physical support and functional control.

### The interaction between vimentin and RRBP1

Among the numerous candidates, we focused on the interaction between vimentin and RRBP1, a positively charged membrane-bound protein found in rough ER (Cui et al. 2012), because: (1) RRBP1 was identified as the top candidates in both mass spectrometry and RNA sequencing examinations. (2) RRBP1 could directly bind viral RNA and play pronounced role during DENV and ZIKV infection (Ooi et al. 2019).

A previous study has identified that RRBP1 could mediate the interaction between ER and MTs via the novel MT-binding and -bundling domain MTB-1 of coiled-coil region of RRBP1 (Ogawa-Goto et al. 2007). Overexpression of MTB-1 induced acetylated MTs and promoted MT bundling (Ogawa-Goto et al. 2007). Moreover, RRBP1 also regulates ER organization and controls axon specification by regulating local MTs remodeling (Farias et al. 2019). Aside from the interplay with MTs, our data demonstrate that RRBP1 colocalizes with ZIKV dsRNA, and knockdown of RRBP1 in wild-type cells reduces ZIKV RNA replication (Fig 6C,H).

Significantly, our results are the first to identify that vimentin can directly bind to RRBP1 and influence its cellular localization and expression level in both mock-infected and ZIKV infected cells. This represents an improved understanding of the interplay between cytoskeletal IFs and ER proteins, especially in the context of virus infection. Of note, RRBP1 depletion has less effect on viral replication than that of vimentin depletion (Fig 6H,I), and RRBP1 expression has no significant influence on the cellular distribution, mRNA and protein expression of vimentin, indicating RRBP1 acts through vimentin during ZIKV infection. Further investigation is needed to determine whether other RNA-binding proteins may cooperate with vimentin to modulate ZIKV replication. Identification of these factors not only benefits the characterization of the biogenesis of ZIKV RCs, but also provides potential candidates for developing broad spectrum compounds that restrict viral replication.

## MATERIA AND METHODS

### Cell culture and virus

Hepatoma Huh7 cells, human osteosarcoma (U2OS) cells and African green monkey kidney epithelial Vero cells were cultured at 37 °C with 5% CO_2_ in Dulbecco’s modified Eagle’s medium (DMEM) (Biological Industries) supplemented with 10% fetal bovine serum (FBS) (Gibco), 1% penicillin and streptomycin. C6/36 cells were cultured in minimum essential medium (MEM) (Gibco) supplemented with 10% FBS and 2% non-essential amino acids (Solarbio, N1250-100) at 28 °C in 5% CO_2_. ZIKV strain SZ01 was used in this study (GneBank: KU866423.2). Virus stocks were prepared by virus amplification in C6/36 cells at a multiplicity of infection (MOI) of 0.1. Virus-containing supernatant medium were harvested from day 4 post infection and stored at −80 □. For lentivirus production, the pLKO.1 shRNA plasmid was transfected into HEK293T cells together with psPAX2 packaging plasmid (Addgene #12260) and pMD2.G envelop plasmid (Addgene #12259) by using FuGENE HD (Promega). Supernatants were collected 48 hours post-transfection, filtered through a 0.45 μm filter to remove the cells debris and stored at −80□. The effectiveness of knockdown of target gene was assessed by qRT-PCR and western blotting.

### Plasmids and transfection

Constructs expressing mCherry-tagged full-length vimentin and GFP-tagged ‘unit length filament’ (ULF) vimentin were kind gifts from John Eriksson (University of Turku and Abo Akademi University, Finland). Plasmids encoding ZIKV envelop was amplified by reverse transcription-PCR (RT-PCR) and cloned into the pEGFP-N1 vector. All constructs were verified by DNA sequencing. The PCR primers used in this study are summarized in Table S3. Transfection of plasmids at indicated concentrations were performed using jetPRIME transfection reagent (#114-15) following the manufacturer’s instructions.

### Vimentin CRISPR knockout cell line generation

As previous, vimentin-knockout cells were generated using CRISPR/Cas9 methods (Jiu et al. 2015) based on pSpCas9 (BB)-2A-GFP vector (Addgene #48138) with two targets. Primers for vimentin target 1 were 5’-CACCGTGGACGTAGTCACGTAGCTC-3’ and 5’-AAACGAGCTACGTGACTACGTCCAC-3’. Primers for vimentin target 2 were 5’-CACCGCAACGACAAAGCCCGCGTCG-3’ and 5’-AAACCGACGCGGGCTTTGTCGTTGC-3’. Transfected cells were detached at 24 h post-transfection and sorted with FACS Aria II (BD Biosciences) using low intensity GFP-expression pass gating, and then cells were plated onto 96-well plate supplemented DMEM containing 20% FBS and 10 mM HEPES with single cell/well. CRISPR clones were cultivated for two weeks prior selecting clones with no discernible vimentin protein expression by western blotting.

### Immunofluorescence microscopy

Cells cultured on glass slides (VWR, #631-0150) were fixed in 4% paraformaldehyde (PFA) for 15 min at room temperature (RT), and permeabilized with 0.1% Triton X-100 in PBS for 5 min. Cells were then blocked in PBS supplemented with 5% bovine serum albumin (BSA) (ABCONE, #A23088). Both primary and fluorescent-conjugated secondary antibodies were applied onto cells and incubated at RT for 1 h. Cells were mounted in DAPI Fluoromount-G reagent (SountherBiotech, 0100-20) and imaged using either Olympus spinSR10 Ixplore spinning disk confocal microscope or GE DeltaVision OMX SR super-resolved structured illumination microscope. The following primary antibodies were used: vimentin rabbit monoclonal D21H3 antibody (dilution 1:100; #5741, Cell Signaling); vimentin chicken polyclonal antibody (dilution 1:1000; #ab24525, Abcam); tubulin mouse monoclonal antibody (dilution 1:200; #4026, Sigma); ribosome-binding protein 1 (RRBP1) rabbit polyclonal antibody (dilution 1:100; #A303-996A, Bethyl Laboratories); Zika virus envelop mouse monoclonal antibody (dilution 1:1000; #1176-46, BioFront); Zika virus NS4B rabbit polyclonal antibody (dilution 1:1000; #GTX133311, Genetex); Zika virus NS1 mouse monoclonal antibody (dilution 1:1000; #1225-06, BioFront); dsRNA monoclonal antibody (dilution 1:500; #J2-1909, SCICONS). The following secondary antibodies were used: Alexa Fluor 488 goat anti-rabbit IgG (H+L) (dilution 1:1000; #A11008, Invitrogen); Alexa Fluor 568 goat anti-rabbit IgG (H+L) (dilution 1:1000; #A11011, Invitrogen); Alexa Fluor 488 goat anti-mouse IgG (H+L) (dilution 1:1000; #A11001, Invitrogen); Alexa Fluor 555 goat anti-mouse IgG (H+L) (dilution 1:1000; #A21422, Invitrogen). F-actin was stained by Alexa Fluor 647 phalloidin (dilution 1:500; #A22287, Invitrogen).

### Live cell imaging

Cells were seeded into 35 mm-diameter glass bottom culture dish (MatTek Corporation) pre-coated by fibronectin (1.5 μg/cm^2^) (Sigma-Aldrich, F2006). Cells were then infected with ZIKV (MOI=5 pfu per cell) for 2 h at 37°C. After removing the inoculum, 2 mL DMEM containing 2% FBS was added for imaging. Alternatively, cells were transfected with ZIKV-E-EGFP for 20 h before imaging. Image series were acquired on an Olympus spinSR10 Ixplore spinning disk confocal microscope using a 100× U plan apochromat high resolution objective with NA=1.5, with time interval of 10 min for 40 h in infection experiments and 13 s for 1.5 h in transfection experiments, respectively. For figure 2C, vimentin knockout cells were co-transfected with ZIKV-E-EGFP and vimentin-mCherry plasmids for 24 h, and image series were acquired on GE DeltaVision widefield microscope using 60× UPlanXApo objective with NA=1.42. Image acquisition was performed at time interval of 30 min for 12 h. All live cell imaging data were further analyzed by Imaris 9.2 (Bitplane, Zurich, Switzerland) and ImageJ software.

### Western blotting

Cells were washed two times with PBS and lysed in RIPA lysis buffer (Beyotime, #P0013B) with protease and phosphatase inhibitors (Beyotime, #P1045). Protein concentration were measured by BCA (Beyotime, #P0010), adjusted with PBS and 6X SDS-sample buffer (Beyotime, #P0015F), and subjected to SDS-PAGE. 5% non-fat milk (BD Difco, #8011939) was used in blocking and PVDF membrane (Millipore, #IPVH00010) was washed by TBST buffer (Tris-buffered saine, 0.1% Tween20). Antibodies were used with the following dilutions in primary antibody dilution buffer (Beyotime, P0023A): vimentin rabbit monoclonal D21H3 antibody (dilution 1:1000; #5741, Cell signaling); phospho-vimentin (Ser39) rabbit antibody (dilution 1:1000; #13614S, CST); phospho-vimentin (Ser56) rabbit antibody (dilution 1:1000; #3877S, CST); phospho-vimentin (Ser83) (D5A2D) rabbit antibody (dilution 1:1000; #12569S, CST); β-tubulin mouse monoclonal antibody (dilution 1:1000; #T4026, Sigma); β-actin mouse monoclonal antibody (dilution 1:1000; #A5441, Sigma); Zika virus envelop mouse monoclonal antibody (dilution 1:5000; #1176-46, BioFront); ribosome-binding protein 1 (RRBP1) rabbit polyclonal antibody (dilution 1:1000; #A303-996A, Bethyl Laboratories); green fluorescent protein (GFP) mouse monoclonal antibody (dilution 1:1000; #G6795, Sigma); p21 Waf1/Cip1 (12D1) rabbit monoclonal antibody (dilution 1:1000; #2947, CST); LC3C (D3O6P) rabbit monoclonal antibody (dilution 1:1000; #14736, CST); GAPDH rabbit monoclonal antibody (dilution 1:5000; #G8795, Sigma). Horseradish peroxidase (HRP)-linked anti-mouse IgG antibody (dilution 1:5000; #7076V, CST) and HRP-linked anti-rabbit IgG antibody (dilution 1:5000; #7074V, CST) were used and chemiluminescence was measured after using western blotting ECL (Tanon, #180-501). The band intensities of blots were measured by ImageJ software. For quantification, the intensities of interested proteins were normalized with the internal control GAPDH, and mock infected cells were set to 1 in each experiment.

### Real-time RT-PCR

Total cellular RNA was extracted by EZ-press RNA Purification Kit (EZBioscience, #B0004DP) according to the manufacturer’s protocols. Total RNA was reverse transcribed by using Color Reverse Transcription Kit (EZBioscience, #A0010CGQ). Real-time RT-PCR was carried out by using 2× Color SYBR Green qPCR Master Mix (ROX2 plus) (EZBioscience, #A0012-R2) in QuantStudio 1 system (Thermo). All readings were normalized to the level of GAPDH. The primers used for real-time RT-PCR are shown in Table S1.

### Plaque assay

Zika virus titers were determined by plaque assay performed on Vero cells. Briefly, Vero cells were seeded into 24 well plates at a density of 1×10^5^ cells/well and washed with pre-warmed phosphate-buffered saline (PBS). Cells were then infected with serial 10-fold dilutions of virus supernatants for 2 h at 37 □ with 5% CO_2_. Inoculum was removed and replaced with DMEM containing 1% carboxymethylcellulose (CMC) (Sigma, #C5678) and 1.5% FBS. After four days post-infection, cells were washed with PBS and fixed with 4% PFA at RT for 1 h, followed by staining with crystal violet (Beyotime, C0121) for 10 min. After rinsing with water, the number of visible plaques was counted, and the virus titers were shown as plaque forming units (PFU) per milliliter.

### Pull-down assay

For preparing the medium, Ni Sepharose 6 Fast Flow (Sigma-Aldrich, GE17-5318-01) was sedimented by centrifugation at 500×g for 5 min, then washed with distilled water and binding buffer (20 mM sodium phosphate, 0.5 M NaCl, 20 mM imidazole, pH 7.4, filtration through a 0.45 μm filter) for twice, and further resuspended with an appropriate volume of binding buffer to make a 50% slurry. For binding the sample, 5 μg of His-tagged recombinant vimentin (Sino Biologicals, #10028-H08B) were incubated with 10 μL of the 50% slurry at 4□ on a shaker with low speed for 3 h. Beads were spun down by centrifugation at 500×g for 5 min and washed with cold binding buffer, then incubated with 1000 µg of filtered whole-cell lysates in 1×Lysis/Binding/Wash buffer (Cell signaling, #11524S) at 4 □ on a rotator with low speed overnight. For elution, Beads were spun down and washed with cold binding buffer for three times, then incubated with cold elution buffer (20 mM sodium phosphate, 0.5 M NaCl, 500 mM imidazole, pH 7.4, filtration through a 0.45 μm filter) at 4 □ on a shaker at low speed overnight. The supernatant was boiled in SDS loading buffer (Beyotime, #P0015F) and subjected to SDS-PAGE, followed by western blotting.

### Mass spectrometry

Cells were lysis with NP40 buffer (50mM pH8.0 Tris-HCl; 150mM NaCl; 0.5%NP40; 1mM EDTA; protease Inhibitor) on ice for 15 min. Scrape cells and harvest the supernatant by centrifuge at 13,000 rpm for 10 min at 4 □. Then proceed immunoprecipitation according to the protocol of Dynabeads Protein G (Thermo Fisher, #10004D). Briefly, incubate Dynabeads Protein G with anti-vimentin antibody (Abcam, #ab137321) with rotation for 30 min at RT. Place the tube with Dynabeads-Ab complex on the magnet to remove the supernatant. Then add the cell supernatant and incubate with rotation for 30 min at RT or overnight at 4 □ to allow antigen to bind to the beads-Ab complex. Wash the Dynabeads-Ab-Ag complex 3 times. Then elute target antigen with SDT buffer (4%SDS; 100mM pH8.0 Tris-HCl; 1mM DTT) for further Mass spectrum analyses by Q Exactive (Thermo Fisher).

### RNA Sequencing and transcriptome analysis

RNA was extracted from U2OS WT or vimentin KO cells after infection with ZIKV (MOI=1) for 24 h, using TRIzol™ Reagent (Invitrogen, #15596026) according to the manufacturer’s instruction. Libraries of RNAs were constructed using Illumina Truseq™ RNA sample prep Kit (Illumina) with the manufacturer’s instruction. Illumina HiSeq 2000 was used to sequence the library and FastQC v0.11.4 (Andrews 2010) was utilized to evaluate the quality of the raw reads. Adapters were removed by Cutadapt v1.16 (Martin 2011) and the same software we used to filter the reads with low quality (Q < 20) or short length (< 25 bp). Hisat2 v 2.1.0 (Kim et al. 2019) was used to map the clean reads on human genome (GRCh38). StringTie v 1.1.3b (Pertea et al. 2016) and GRCh38 GTF file (version 92) were used to obtain the counts of genes and edgeR v 3.24 (McCarthy et al. 2012, Robinson et al. 2010) was used to perform the analysis of differential gene expression analysis. FPKM of each gene was calculated and different expression genes which |log2FoldChange| > 1 and adjust P-value < 0.05 were selected. Volcano plot and Heat map were generated using TBtools (Chen et al. 2020).

### GO enrichment analysis

Gene ontology analysis was performed by using the Database for Annotation, Visualization and Integrated Discovery (DAVID) Bioinformatics Resources 6.8 (https://david.ncifcrf.gov/home.jsp) (Huang da et al. 2009). The molecular function, biological process and cellular component of tested genes were analyzed. Significance was defined as adjust P-value (Padj) less than 0.05. The results were presented with bubble plots drawn by R with gglpot2 package (Wickham 2016).

### Transmission electron microscopy

Cells were seeded on ACLAR films treated with poly-lysine (100 ug/mL), infected with ZIKV (MOI=5) for 24 h, and fixed in 2.5% glutaraldehyde in PBS with a pH of 7.4 at room temperature for 1 hour. Samples were then post-fixed with 1% osmium tetroxide in 0.1 mol/L sodium cacodylate for 1.5 h, and stained with 2% (w/v) uranyl acetate in double distilled water for 50 min to increase the contrast. After washing and dehydration in a graded series of acetone, samples were embedded in Embed 812 resin. The embedded samples were sliced into 70 nm sections using a Leica ultramicrotome EM UC6 (Leica, Germany) and sections were collected on the EM grids. Sample grids were imaged under a spirit transmission electron microscope (FEI Company, The Netherlands) operating at 100 Kv.

### Statistical analysis

Statistical analysis was performed by unpaired Student’s *t* test, one-way analysis of variance or two-way ANOVA using the GraphPad Prism v8 software and p values were indicated by *p<0.05, or **p<0.01, or ***p<0.001. The histogram data were presented as mean ± SEM.

## ACKNOWLEDGMENTS

We would like to thank John E. Eriksson (University of Turku and Åbo Akademi University, Finland) for providing us the vimentin plasmids, Lan Bao (Chinese Academy of Science, China) and Xueliang Zhu (Chinese Academy of Science, China) for insightful discussion.

## AUTHOR CONTRIBUTIONS

Y.J. and X.J. designed and supervised the study. Y.Z. carried out experiments and interpretation of the data. Y.X. and F.F. performed the electronic microscopy. J.C. and Y.L. performed the RNA-Seq experiment. S.Z. performed the Mass Spectrometry experiment. Y.J., X.J. and Y.Z. wrote the manuscript with contributions from all other authors.

