## supplemental figure legends for "Host cytoskeletal vimentin serves as a structural organizer and an RNA-binding protein regulator to facilitate Zika viral replication"

**Figure S1 Vimentin filaments, but not microtubules and actin filaments, are dramatically reshaped upon ZIKV infection. (A)** Time course of immunoblot of total protein levels and the phosphorylated levels of vimentin in non-infected (MOCK) and ZIKV-infected cells (MOI=0.5). GAPDH is used to verify equal sample loading. **(B)** Immunoblots of distinct vimentin cellular fractions (membrane-associated, cytoplasmic, cytoskeleton-associated, and nuclear-associated) in MOCK and ZIKV-infected cells (MOI=1). **(C)** Immunofluorescence of microtubules and actin filaments network in mock and ZIKV-infected cells (MOI=5). **(D)** Effect of microtubule destruction on the localization of vimentin and viral proteins upon ZIKV infection (MOI=5). Dynein inhibitor Cilibrevin D (20 μM) was added at 2 hpi for 24 h, nocodazole (10 μM) was added for at 12 hpi for 12 h. Scale bar 15 μm. **(E)** Cells infected with ZIKV for 48 h (MOI=5) were immunostained for vimentin and NS1 or NS4B. Scale bar 15 μm. **(F)** Cells transfected with pEGFP-N1 vector were immunostained with vimentin and actin. Scale bar 15 μm. **(G)** Transmission electronic microscopy images of 70 nm thin sections of resin-embedded cells infected with ZIKV (MOI=5). White dotted box in the left panel indicates the magnified area shown in the corresponding color image on the right. Ve, viral induced invaginated vesicles, blue; Vi, ZIKV virions, green; zER, zippered ER, purple. Scale bar 5 μm in the cell images, 100 nm in the magnified images.

**Figure S2 Vimentin knockout leads to replication compartments disruption and thus defective ZIKV infection. (A)** Western blot verified the vimentin KO and full-length rescue efficiency in U2OS cells. **(B)** Quantification of the viability rates of U2OS WT and vimentin knockout (VIM KO) cells. **(C)** Cell growth curves in WT and VIM KO U2OS cells determined by cell numbers. **(D)** Western blot verified the VIM KO Huh7 cells. **(E)** Cell growth curves in WT and VIM KO Huh7 cells determined by cell numbers. **(F)** ZIKV Env protein level in infected WT and VIM KO Huh7 cells (MOI=0.1) were measured by western blotting. **(G)** Intracellular ZIKV RNA levels in infected WT and VIM KO Huh7 cells (MOI=0.1) were measured by qRT-PCR. The data are from three independent experiments. **(H)** ZIKV RNA levels in the supernatant of infected WT and VIM KO Huh7 cells (MOI=0.1) were measured by qRT-PCR. The data are from three independent experiments. **(I)** WT and VIM KO U2OS cells were infected with ZIKV (MOI=5) for 24 h. Cell were fixed and immunostained to visualize ZIKV NS1/NS4B, vimentin, actin and nucleus. Scale bar 15 μm. **(J)** Quantification of the percentages of cells with segregated ZIKV Env/NS1/NS4B in WT and VIM KO U2OS cells. Each point represents an independent experiment. GAPDH were used in all western blots to verify equal sample loading. Error bars indicated means ± SEM.

**Figure S3 RNA-seq analysis. (A-B)** GO analysis of significantly down-regulated genes between ZIKV infected WT and vimentin KO U2OS cells. (MOI=1, 24 hpi). The color of the bubbles displayed from red to blue indicated the descending order of -log10(Padj). The sizes of the bubbles are displayed from small to large in ascending order of gene counts. The x and y axis represent the gene ratio and the GO terms, respectively. (A) All significant GO terms in cellular component (A) and molecular function (B) are shown in the bubble chart. **(C)** Volcano plot highlighting differentially expressed genes between ZIKV infected WT and vimentin KO U2OS cells (MOI=1, 24 hpi). Color dots denote up- and down- regulation of individual candidates indicated as red and blue, respectively.

**Figure S4 Vimentin regulates ZIKV infection by controlling of ER-located RRBP1. (A)** Protein levels of RRBP1, viral Env and vimentin in WT and vimentin knockout (VIM KO) U2OS cells analyzed by western blots. **(B)** Quantifications of the band intensities of RRBP1 in western blots upon ZIKV infection in WT and VIM KO U2OS cells. Band intensity of RRBP1 were normalized to GAPDH. Each point represents an independent experiment. ***P*<0.01 (unpaired *t*-test). **(C)** Quantifications of the mRNA levels of RRBP1 in WT and VIM KO U2OS cells measured by qRT-PCR. Each point represents an independent experiment; ****P*<0.001 (unpaired t-test). **(D)** Quantification of colocalization between RRBP1 and vimentin in U2OS non-infected and ZIKV-infected cells was performed using Pearson’s coefficients. Each point represents a single cell; ****P*<0.001 (unpaired *t*-test). **(E)** Relative mRNA levels of RRBP1 in WT, VIM KO, RRBP1 KD and RRBP1 KD; VIM KO cells. Each point represents an independent experiment. **P*<0.05, ***P*<0.01, ****P*<0.001 (two-way ANOVA) **(F)** Relative mRNA levels of vimentin in WT, VIM KO, RRBP1 KD and RRBP1 KD; VIM KO cells. Each point represents an independent experiment. **P*<0.05 (two-way ANOVA). Error bars indicated means ± SEM.

**Video S1 related to Figure 1D (upper panel)** Time-lapse video of vimentin-mCherry stably expressing U2OS cells in normal culture condition. The display rate is 16 frames per second. Scale bar 15 µm.

**Video S2 related to Figure 1D (lower panel)** Time-lapse video of vimentin-mCherry stably expressing U2OS cells infected with ZIKV (MOI=5). The display rate is 16 frames per second. Scale bar 15 µm.

**Video S3 related to Figure 2A** Time-lapse video of vimentin-mCherry stably expressing U2OS cells transfected with ZIKV Env-EGFP. The display rate is 10 frames per second. Scale bar 15 µm.

**Video S4 related to Figure 2C** Time-lapse video of U2OS vimentin knockout cells co-transfected with ZIKV-Env-EGFP and vimentin-mCherry plasmids. The display rate is 4 frames per second. Scale bar 5 µm.

**Table S1 Summary of ER-located proteins interacting with vimentin by Mass Spectrometry**

| Protein ID | Protein name | Gene name | Coverage (%) |
| --- | --- | --- | --- |
| P62820 | Ras-related protein Rab-1A | RAB1A | 48.78 |
| Q07065 | Cytoskeleton-associated protein 4 | CKAP4 | 48.34 |
| P62857 | 40S ribosomal protein S28 | RPS28 | 46.38 |
| P50402 | Emerin | EMD | 41.73 |
| Q02878 | 60S ribosomal protein L6 | RPL6 | 38.89 |
| O14880 | Microsomal glutathione S-transferase 3 | MGST3 | 37.5 |
| J3QQY2 | Calcium load-activated calcium channel | TMCO1 | 35.58 |
| P61247 | 40S ribosomal protein S3a | RPS3A | 35.23 |
| Q15717 | ELAV-like protein 1 | ELAVL1 | 34.36 |
| Q15050 | Ribosome biogenesis regulatory protein homolog | RRS1 | 32.05 |
| O43390 | Heterogeneous nuclear ribonucleoprotein R | HNRNPR | 31.6 |
| Q03135 | Caveolin-1 | CAV1 | 30.9 |
| A7BI36 | p180/ribosome receptor | RRBP1 | 30.58 |
| P11021 | Endoplasmic reticulum chaperone BiP | HSPA5 | 30.28 |

**Table S2 Summary of identified RRBP1 peptides interacting with vimentin by Mass Spectrometry**

| Peptides | Alignment site |
| --- | --- |
| QLLLESQSQLDAAK | 1228-1241 |
| ETSYEEALANQR | 18-46 |
| TLVSTVGSMVFNEGEAQR | 652-669 |
| NTDVAQSPEAPK | 609-620 |
| QSDELALVR | 1247-1255 |
| SVEEEEQVWR | 1148-1157 |
| DAQDVQASQAEADQQQTR | 971-988 |
| QVLQLQASHR | 865-874 |
| KLQEQLEK | 1390-1397 |
| LKGELESSDQVR | 1184-1195 |
| LQQENSILR | 798-806 |
| ESEEALQK | 875-882 |
| EVPMVVVPPVGAK | 115-120 |
| HLEEIVEK | 1176-1183 |
| EQEITAVQAR | 752-761 |
| VDTTPNQGK | 170-176 |
| EVQQLQGK | 772-779 |
| LQSSEAEVR | 922-930 |
| LHSLTQAK | 1042-1049 |
| LIEILSEK | 670-677 |
| HPPAPAEPSSDLASK | 1099-1113 |
| AQEQQQQMAELHSK | 908-921 |
| KAEGTPNQGK | 632-666 |
| LLATEQEDAAVAK | 708-720 |
| SILAETEGMLR | 1133-1143 |
| VGAAEEELQK | 1160-1169 |
| LKELESQVSGLEK | 989-1001 |
| LTSDLGR | 1359-1365 |

**Table S3 Sequences of primers used in this work**

|  |  | Primers | Sequence (5’-3’) |
| --- | --- | --- | --- |
| PCR primer | Envelop | Forward | CGAAGCTTATGATCAGGTGCATAGGAGTCAGCA |
|  |  | Reverse | CGGGATCCCGAGCAGAGACGGCTGTGGATAAG |
| qPCR primer | ZIKV | Forward | CAACCACAGCAAGCGGAAG |
|  |  | Reverse | AAGTGATCCATGTGATCAGTTGATCC |
|  | RRBP1 | Forward | TTCAACGAGGGCGAGGCCCAG |
|  |  | Reverse | CGTGCCTGCACAGCCGTGATCT |
|  | GAPDH | Forward | GCATCCTGCACCACCAACTG |
|  |  | Reverse | GCCTGCTTCACCACCTTCTT |
|  | Vimentin | Forward | GACCTTGAACGCAAAGTGGAATC |
|  |  | Reverse | GTGAGGTCAGGCTTGGAAACATC |
