## supplemental figure 1-4 for "Host cytoskeletal vimentin serves as a structural organizer and an RNA-binding protein regulator to facilitate Zika viral replication"

#### Figure S1 related to Figure 1 and Figure 2

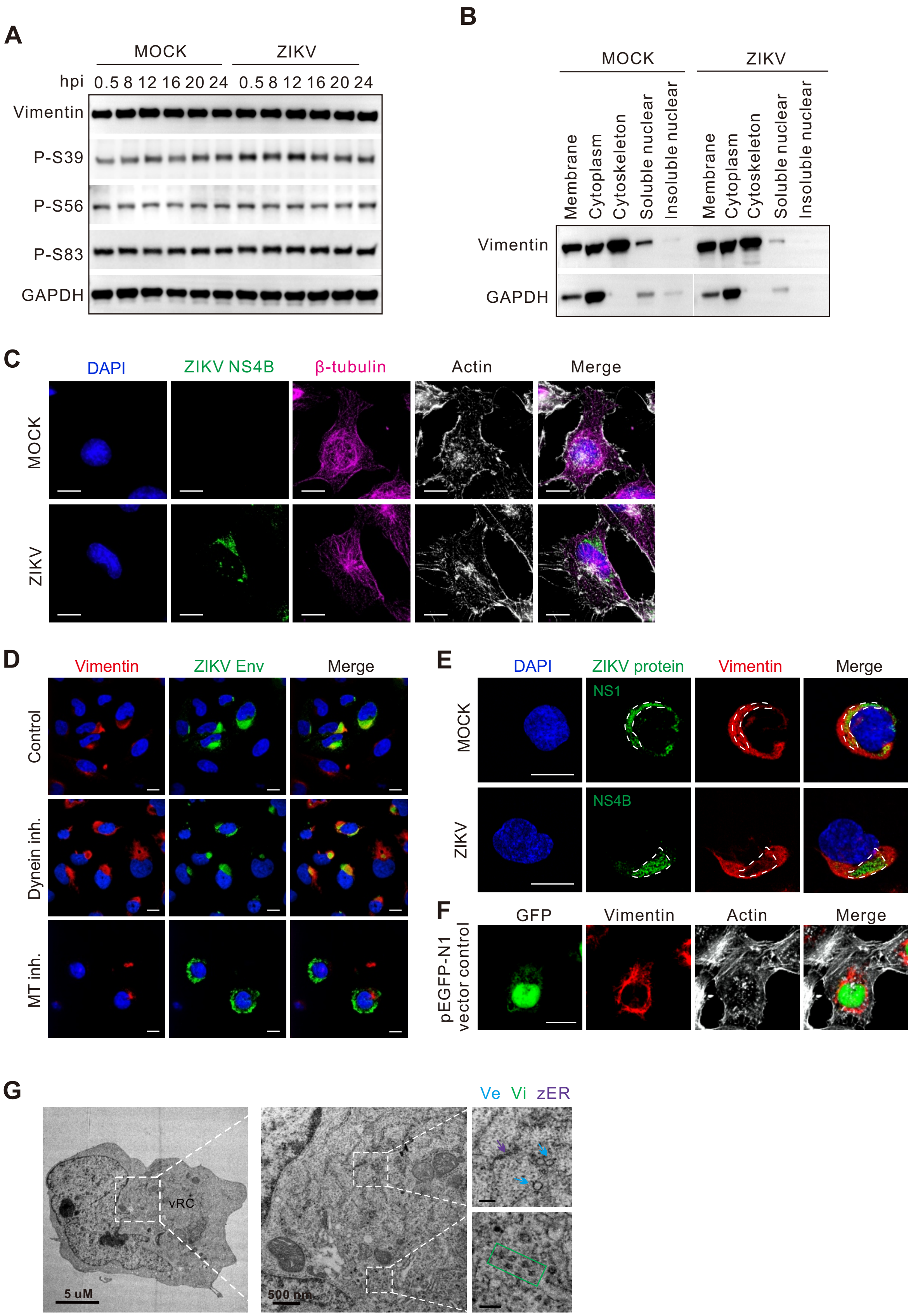

#### Figure S2 related to Figure 3

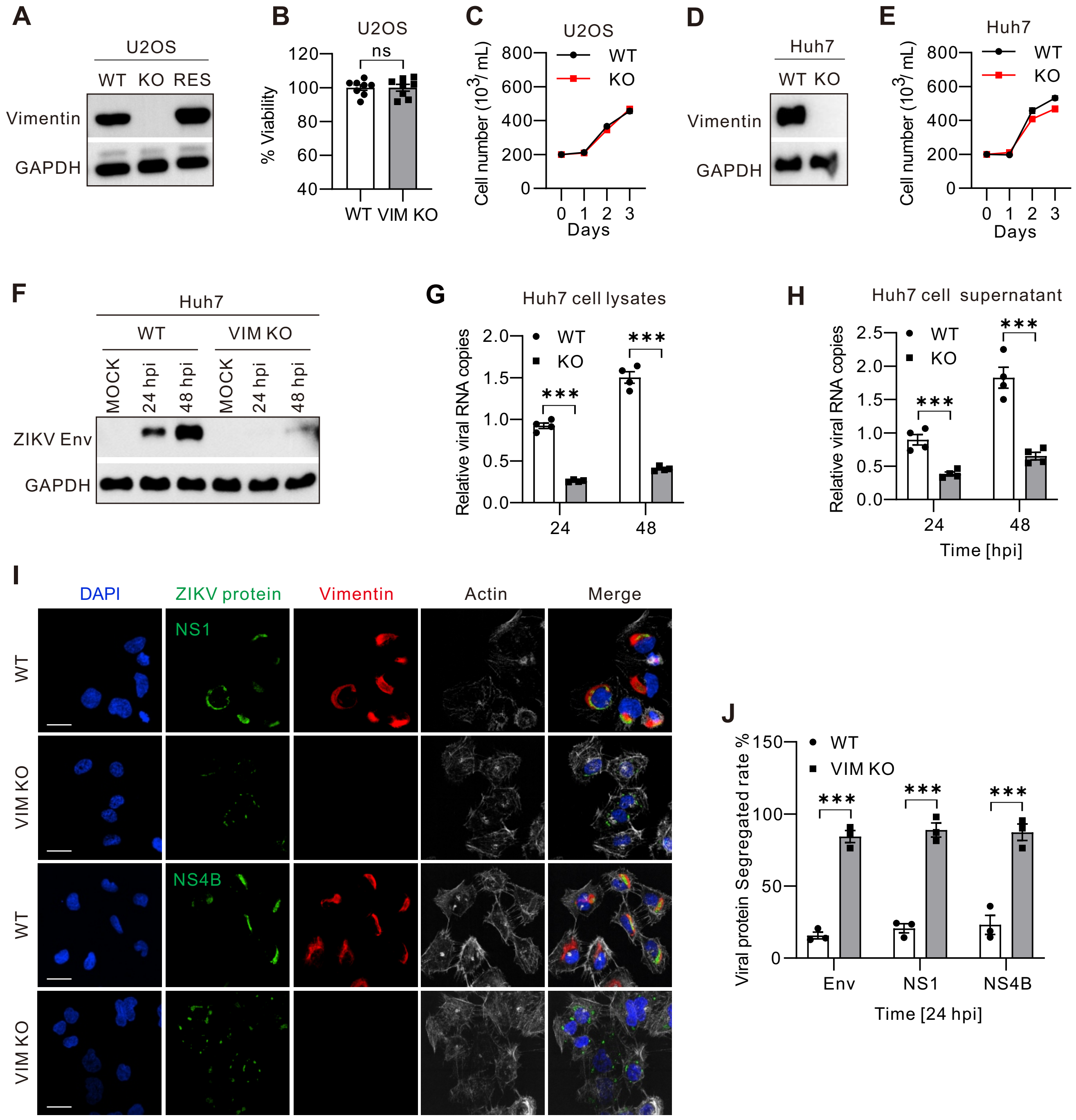

### Figure S3 related to Figure 5

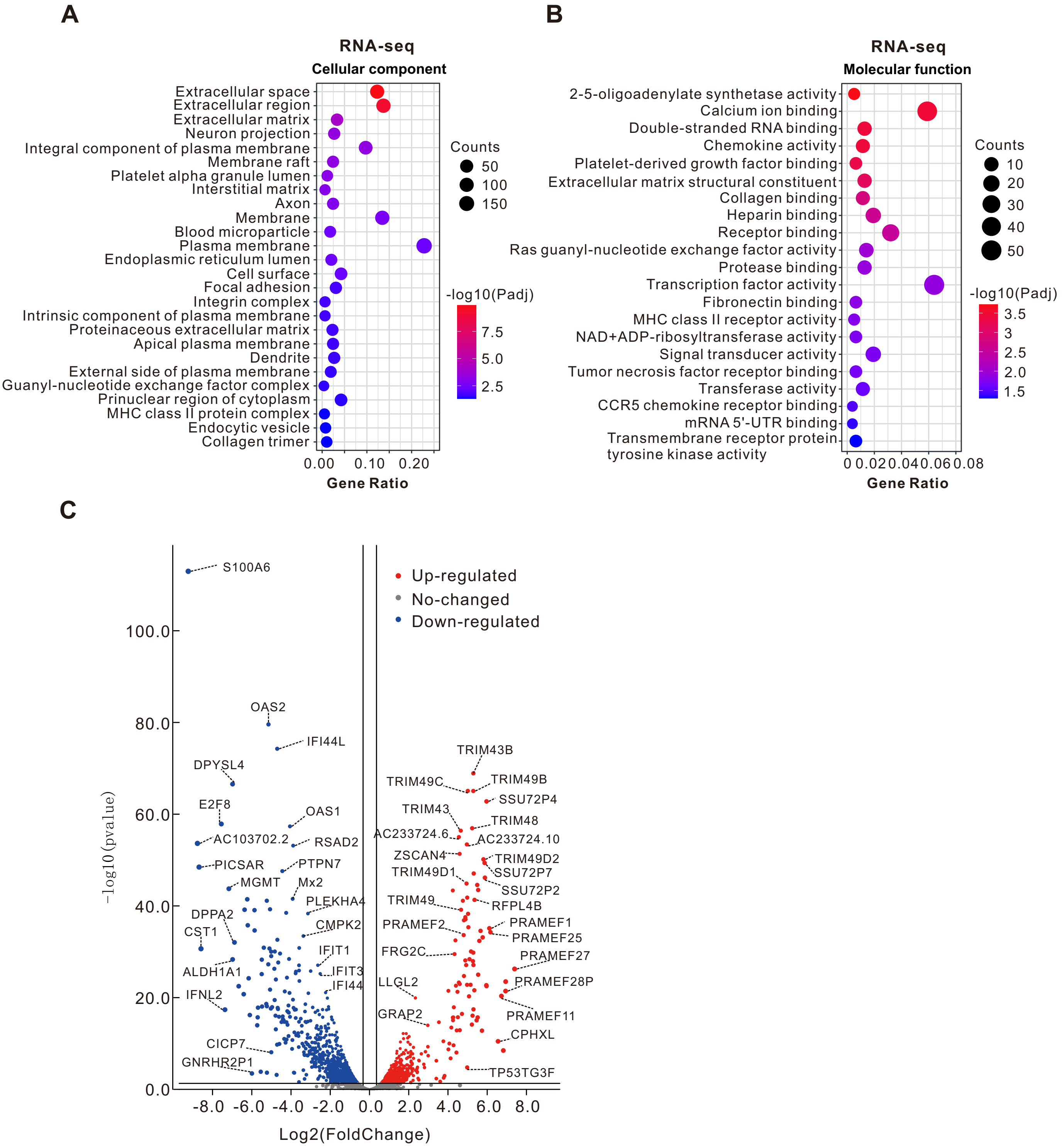

#### Figure S4 related to Figure 6

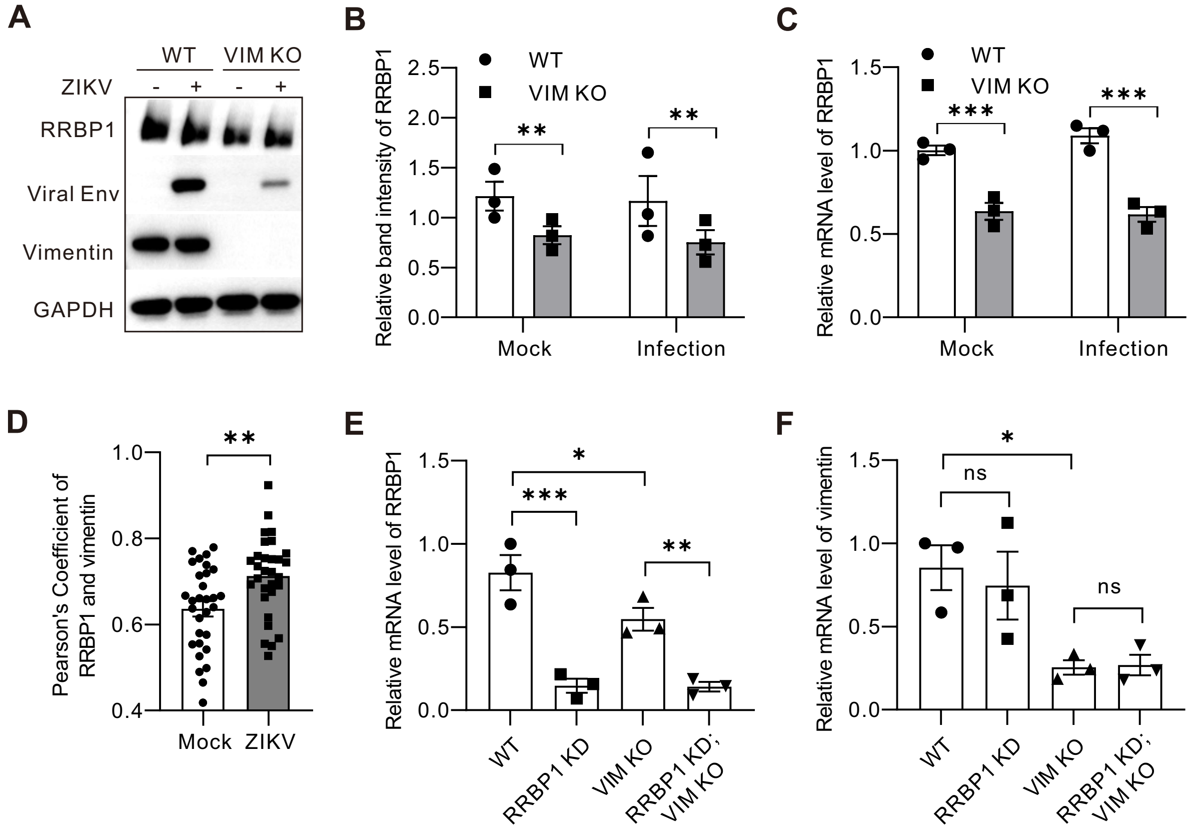
